# A Single Cell Genomics Atlas of the *Drosophila* Larval Eye Reveals Distinct Developmental Timelines and Novel Markers for All Photoreceptor Subtypes

**DOI:** 10.1101/2023.02.14.528565

**Authors:** Komal Kumar Bollepogu Raja, Kelvin Yeung, Yoon-Kyung Shim, Yumei Li, Rui Chen, Graeme Mardon

## Abstract

The *Drosophila* eye is a powerful model system to study principles of cell differentiation, proliferation, survival and morphogenesis. However, a high-resolution single cell genomics resource that accurately captures all major cell types of the larval eye disc and their spatiotemporal relationships is lacking. Here, we report transcriptomic and chromatin accessibility data for all known cell types in the developing eye. Photoreceptors appear as streams of cells that represent dynamic developmental timelines. Photoreceptor subtypes are transcriptionally distinct when they begin to differentiate, but then converge upon a common transcriptome just 24 hours later. We identify novel cell type-specific marker genes, enhancers and potential regulators, as well as genes with distinct R3 or R4 photoreceptor specific expression. Finally, we observe that photoreceptor chromatin accessibility is more permissive than non-neuronal lens-secreting cone cells, which show a more restrictive chromatin profile. This single cell genomics atlas will greatly empower the *Drosophila* eye as a model system.

## Introduction

Biological tissues with complex mixtures of cellular identities, as well as tissues that show rapidly changing temporal patterns of gene expression often require investigations at single cell resolution for deep mechanistic understanding. Recent technological advances have enabled profiling of transcriptomics, epigenomics, and chromatin configuration from complex tissues at single cell resolution. Single cell data provide critical insights into genetic networks underlying development and disease, and have transformed our understanding of biological processes.

Indeed, single cell molecular atlases of tissues and organs from many species, including numerous model organisms^1–5^, have been recently published and are now an essential resource for understanding conserved mechanisms underlying cell fate specification, differentiation and proliferation, as well as the discovery of previously unknown cell types. One of the most commonly used model organisms is *Drosophila*, which has been extensively employed for more than a century to study genetics, development, neuroscience, aging, disease, and many other processes^6–9^. The *Drosophila* eye has been of particular utility since it is easily assayed as an externally visible organ in living animals and has served as a powerful genetic screening tool for decades. Importantly, many genes involved in retinal determination, such as *Pax6* (*eyeless* in flies), are highly conserved between humans and flies and are required for eye development in both species^10–12^. Therefore, it is of great importance to establish high-resolution single cell atlases for different *Drosophila* tissues, and although this has been accomplished for the brain, olfactory projection neurons, and several other tissues, such data has not been reported for the *Drosophila* eye^1, 13–15^.

The adult *Drosophila* compound eye is made of ∼750 repeating hexagonal units called ommatidia. Each ommatidium has eight photoreceptor cells (PRs), four non-neuronal lens-secreting ’cone’ cells, and six pigment cells. The eye develops from a neuroepithelial sac called the eye imaginal disc during larval and pupal stages. A wave of differentiation called the morphogenetic furrow (MF) begins at the posterior margin of the early larval eye disc and moves anteriorly, leaving differentiating cells behind it. Posterior to the MF, the R8 PR differentiates first and is the founder cell for each ommatidium. This is followed by the progressive recruitment and differentiation of three pairs of PRs (R2/5, R3/4, and R1/6). The R7 PR and four cone cells are added to each ommatidial cluster during late larval stages (Fig. 1B, B’). Pigment cells differentiate during pupal development. Furthermore, a new column of ommatidia emerges from the MF every 1.5 hrs such that each column is developmentally more mature than the one immediately anterior to it. Therefore, cells in the larval eye disc are arranged as a developmental space-time continuum. More mature differentiating cells are found toward the posterior of the eye disc while less mature uncommitted progenitor cells are located anteriorly, poised to undergo differentiation. This progressive, temporal component is a unique aspect of the larval eye compared to most other *Drosophila* tissues. Moreover, ommatidia in the dorsal and ventral halves rotate 90° in opposite directions, resulting in dorsal and ventral ommatidia becoming mirror images of each other (Fig. 1B’). Ommatidial rotation is tightly linked to R3 and R4 differentiation and the interaction between the Frizzled/Dishevelled (Fz/Dsh) and Notch (N) signaling pathways is essential for this process^16, 17^. Although several components of these pathways are known, genes that show R3- or R4-specific patterns of expression have not yet been identified to the best of our knowledge. Moreover, even though cell type differentiation, survival, ommatidial polarity, and other developmental processes in the larval eye are highly synchronized and regulated by conserved signaling pathways and gene regulatory networks, much remains to be deciphered. Therefore, single cell genomics resources for the *Drosophila* larval eye that reflect the repeating, highly ordered, and spatiotemporal properties of this tissue will be of great value.

**Fig. 1:**
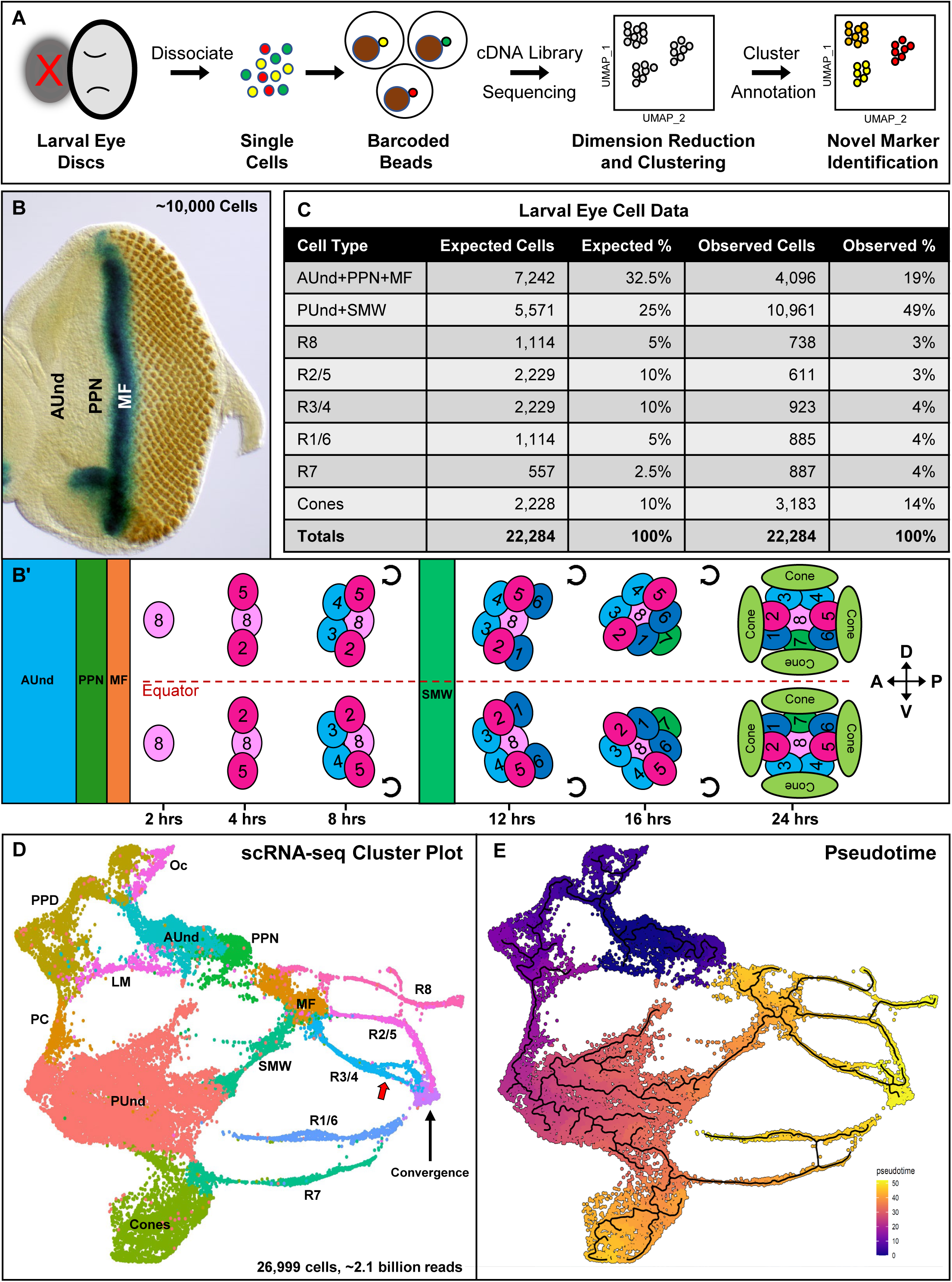
Single cell RNA sequencing of the developing late larval *Drosophila* eye disc reveals all expected cell identities. **A.** Schematic of single cell RNA sequencing data generation and analyses. The red ’X’ indicates that the antennal disc was discarded during dissection. **B.** Larval eye disc carrying a *dpp-lacZ* reporter construct were stained with the neuronal marker Elav (rust-colored dots) and β- galactosidase to visualize *dpp* expression (blue). *dpp-lacZ* is expressed in the MF. **B’.** Schematic depicting the arrangement of cell types according to their developmental age in the larval eye disc. Each cluster of cells represents one ommatidium; a total of 12 ommatidia are depicted. Anterior/left to the MF, AUnd and PPN cells are undifferentiated and poised to begin differentiation. Posterior/right to the MF, the R8 photoreceptor differentiates first, followed by R2/5 and R3/4. All undifferentiated cells then undergo one more round of cell division termed the second mitotic wave (SMW). Following the SMW, R1/6, R7 and cone cells are then recruited. The equator is shown as a red dashed line. Dorsal and ventral ommatidia rotate in opposite directions and exhibit chirality. The direction of rotation is shown as semi- circle arrows. The approximate timing of events is shown below the schematic where t=0 is when cells first exit the MF. **C.** The expected and observed numbers and percentages of cells for each cell identity are shown. Non-eye disc clusters (PPD, PC, LM and Oc) were excluded and only major cell type numbers were used for percentage calculations. **D.** UMAP cluster plot generated from ∼27,000 late larval eye disc cells (after filtering to remove low quality and non-eye disc cells) shows all expected cell identities. Clusters appear in a temporal progression from left to right and their arrangement closely correlates with the physical eye disc. The red arrow points to the R3 and R4 ’split’ in the R3/4 stream. **E.** UMAP cluster plot showing pseudotime analysis of scRNA-seq data. Dark blue indicates early and yellow denotes late pseudotime. The UMAP plot shows the expected pseudotime profile with AUnd cells showing early pseudotime while the Convergence cluster is late. AUnd: Anterior Undifferentiated, PPN: Preproneural, MF: Morphogenetic Furrow, SMW: Second Mitotic Wave, PUnd: Posterior Undifferentiated, PC: Posterior Cuboidal Margin Peripodium, LM: Lateral Margin Peripodium, PPD: Anterior Peripodial, and Oc: Ocelli.

Although single cell data from the *Drosophila* larval eye have been previously reported^18–20^, the data presented in this report differ substantially. Here, we present single cell transcriptomic and chromatin accessibility data from late larval eye discs that comprise a deep representation of all known cell types in this tissue. Our data show that all cell clusters appear in a temporal progression resembling the arrangement of cells in the physical eye disc. Photoreceptor cell clusters appear as distinct streams of cells that emerge from undifferentiated clusters but later converge upon a common transcriptome. We identify dozens of novel cell type-specific markers and enhancers, including genes that distinguish the R3 and R4 photoreceptors, and also present *in vivo* validation of many such markers. Finally, we show that the chromatin accessibility of PRs is clearly distinct from that of cones. Our high-resolution data provides a valuable platform for investigating *Drosophila* eye development, as well as greatly aiding research groups that use the eye disc as a model system.

## Results

### A robust single cell transcriptomic atlas of the developing *Drosophila* eye

We performed single cell RNA sequencing (scRNA-seq) on two biological replicates of late larval eye discs using the 10x Genomics Chromium platform (Fig. 1A). After removing dead or dying cells, multiplets, and non-retinal cells (e.g., brain and glia), this data set contains 26,999 cells with a sequencing depth of ∼2.1 billion reads and 2,173 median genes per cell (detailed metrics are provided in Supplementary Fig. 1A). Importantly, the UMAP plot shows distinct clusters corresponding to all expected cell identities in the larval eye disc (Fig. 1D). Since a single late larval eye disc consists of ∼10,000 cells, the cellular coverage of our data is more than twice the number of cells in the physical eye disc. In addition, as there are only about a dozen different cell identities at this stage, we expect that each cell type is well represented in our dataset and this is indeed the case (Fig. 1C).

### Mapping cell identities to clusters using known marker gene expression

We assigned cell identities in the UMAP cluster plot using marker genes known to be expressed in specific cell types at this stage. For instance, we identified the preproneural (PPN) cluster using *hairy* (*h*), which is expressed in the PPN and negatively regulates the progression of the MF^21^ (Fig. 2A,J). The PPN is a subset of the anterior undifferentiated (AUnd) region that is marked by the expression of *Optix*^22^. While *Optix* is expressed in all AUnd cells (including the PPN, Fig. 2B,J), *h* expression is restricted to the PPN cluster. The morphogen *decapentaplegic* (*dpp*) is required for the propagation of the MF and its expression marks the MF and lateral margins (LM) of the eye disc^23, 24^ (Fig. 1D and Fig. 2C,J).

**Fig. 2:**
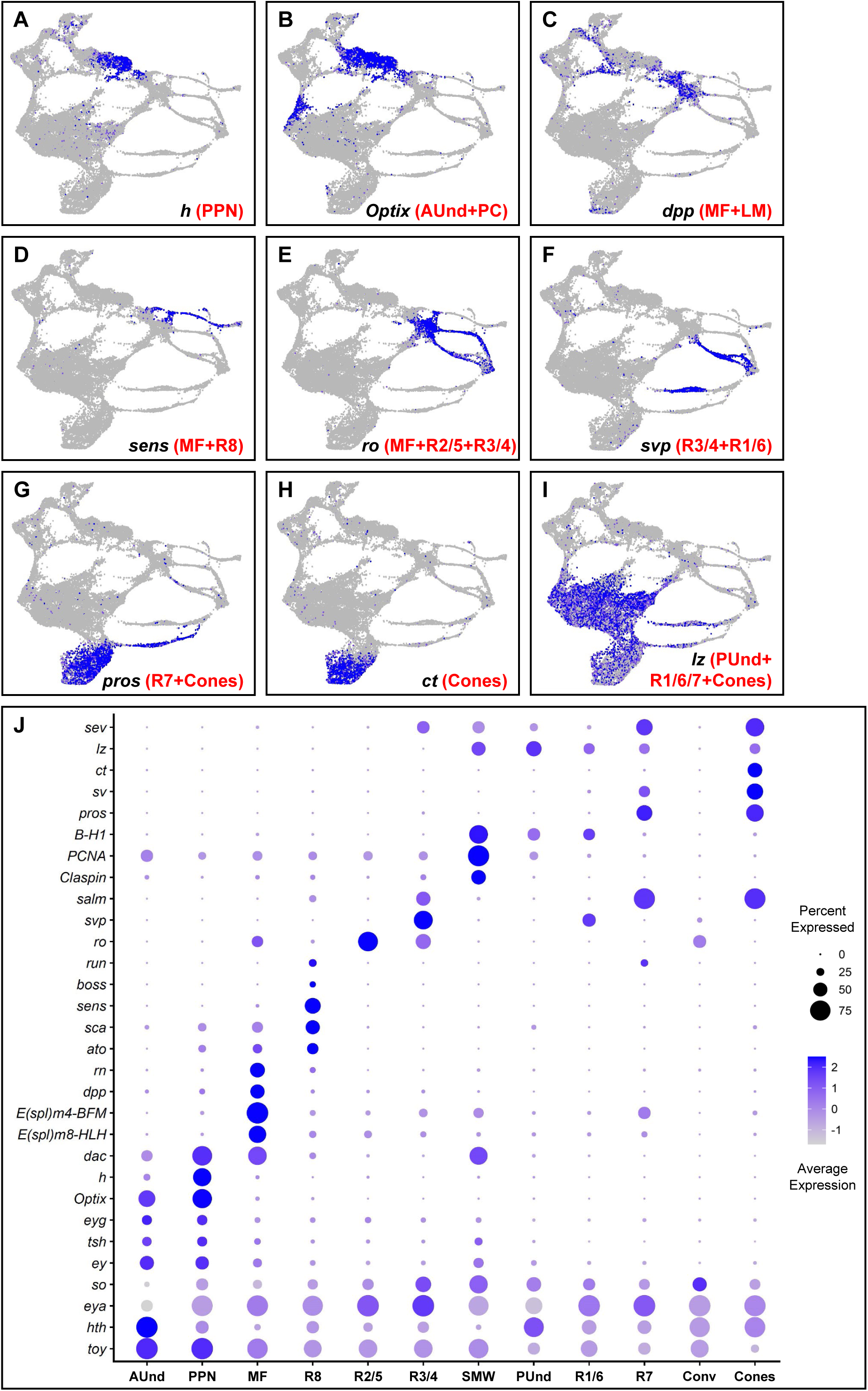
Validation of cluster annotation using known markers. A-I. FeaturePlots showing the expression of marker genes (shown in blue) that were used to identify and annotate clusters. The intensity of blue is proportional to the log-normalized expression levels. **A.** *hairy* (*h*) expression is confined to PPN cells. **B.** *Optix* expression in AUnd cells. **C.** *dpp* expression in the MF and lateral margins. **D.** *senseless* (*sens*) expression in the late MF and R8 cluster. **E.** *rough* (*ro*) is expressed in the MF, R2/5, and R3/4. **F.** *seven up* (*svp*) is expressed in R3/4 and R1/6. **G.** *prospero* (*pros*) is expressed in R7 and cone cells. **H.** *cut* (*ct*) is expressed in cones. **I.** *lozenge* (*lz*) is expressed in PUnd cells and R1/6/7. **J.** Dot plot showing the expression of known marker genes. The intensity of blue denotes the average expression level of each gene in each cluster. The size of the circle is proportional to the percentage of cells in each cluster that express the gene. Known marker genes show specific and expected patterns of expression, supporting the assignment of clusters as shown in Figure 1D.

We identified three cell populations that correspond to the first PR subtypes to differentiate (R8, R2/5 and R3/4; Fig. 1B’). These PR clusters appear as thin streams of cells emerging from the MF. One of the streams shows *atonal* (*ato*), *senseless* (*sens*), and *bride of sevenless* (*boss*) (Fig. 2D,J), which are specifically expressed in R8^25–27^ and are required for R8 differentiation, development, and function^27, 28^. We therefore identify this cluster as R8. We also identified the R2/5 and R3/4 PR clusters using *rough* (*ro*) and *sevenup* (*svp*) as known markers. *ro* is expressed in R2/5 and R3/4^29^ (Fig. 2E,J), whereas *sevenup* (*svp*) is expressed in R3/4 and R1/6^30^ (Fig. 2F,J). Our FeaturePlots show that *ro* is expressed in two PR streams and one of these clusters also coexpresses *svp*. We therefore identify the latter cluster as R3/4 while the other cluster that specifically expresses *ro* but not *svp* represents R2/5. The cluster that expresses *svp* without *ro* expression is annotated as R1/6. We also observe that *Bar*-*H1* (Fig. 2J) and *Bar-H2* (*B-H1* and *B-H2*), known markers of R1/6^31^, are expressed in the R1/6 cluster as well as posterior undifferentiated (PUnd) cells.

The transcription factor *prospero* (*pros*) is expressed in R7 PRs and in non-neuronal cone cells which secrete the lens^32^ (Fig. 2G,J). We observe that *pros* is expressed in two clusters, one of which also expresses *cut* (*ct*), a known cone cell marker^33^. We therefore assigned *pros*- expressing cells that do not express *ct* as R7 and cells that express *ct* and *pros* as cone cells (Fig. 2H,J). After differentiation of R8, R2/5 and R3/4, all remaining undifferentiated cells undergo another round of division known as the second mitotic wave (SMW). R1/6, R7 and cones differentiate from cells following the SMW. A cluster separating the MF and PUnd clusters shows expression of the known cell cycle markers *Claspin* and *Proliferating cell nuclear antigen* (*PCNA*)^18^ and we therefore identify this as the second mitotic wave (SMW) cell cluster (SMW, Fig. 2J). As expected, our UMAP plot shows that the R1/6, R7 and cone cell clusters do not emerge from the MF, but appear from PUnd cells that are adjacent to the SMW cluster. In addition, we observe that most PR subtypes appear to merge into a common cluster, which we named the ’Convergence’ cluster and is discussed in greater detail below.

We identified PUnd cells using *lozenge* (*lz*), which is expressed in PUnd cells as well as in R1/6/7 and cones^34, 35^ (Fig. 2I,J). We also identified cell clusters corresponding to the peripodial membrane (PPD), posterior margin cuboidal cells (PC) and ocelli (Oc) using known marker gene expression (Fig. 1D and Supplementary Fig. 1B-I). The expression patterns of many other genes observed in our data, as revealed by FeaturePlots such as those shown in Fig. 2A-I, are also highly consistent with published studies (Fig. 2J and Supplementary Fig. 1B). Taken together, these results show that our scRNA-seq data accurately represents the endogenous mRNA distribution of all genes examined in the larval eye disc.

Remarkably, the pattern of cell clusters in our dataset closely correlates with the physical eye disc. Our cluster plot shows that AUnd cells and differentiated PR clusters are separated by the MF and PPN clusters in the expected order. Moreover, PR clusters appear as thin streams of cells and the cone cluster is distinct from R7 and PUnd cells. A cluster of SMW cells separates the MF and PUnd (Fig. 1D). Thus, the posterior, differentiated part of the eye disc corresponds to the right side of the cluster plot, while anterior, developmentally less mature cells are toward the left (Fig. 1B,D). Moreover, within each differentiated PR cluster, cells are arranged as a temporal and developmental progression. As expected, trajectory analyses also show that pseudotime progresses from the less mature AUnd cluster to the distal tips of the PRs (Fig. 1E). These data therefore afford a virtual transcriptomic representation of the physical eye disc that is accurate in both space and time. Taken together, our single cell transcriptomic atlas comprises all expected cell types present in the late larval eye disc and all clusters are unambiguously identified and distinct from one another.

### Identification of cell type-specific novel markers

We performed differential gene expression analyses on all cell clusters and identified novel cell type-specific markers for each. We visualized the expression of marker genes and ranked them into four major classes to find markers based on cell type specificity (Supplementary Table1). Class 1 includes genes showing strong and specific expression in one or two eye disc clusters; expression in non-eye disc clusters such as the PPD, PC, LM and Oc may also be present. Descriptions of all four expression classes and the total number of markers observed in each class for each cell cluster are shown in Supplementary Table 1.

Differential gene expression analyses of the R8 stream reveal 1,306 marker genes (Supplementary Table 1), including seven Class 1 markers. One such example is *CG42458* (Fig. 3B,G); several others are shown in Supplementary Fig. 3A-C. To test if *CG42458* is indeed specifically expressed in R8 cells *in vivo*, we used a *CG42458-Trojan-Gal4* (*T2A-Gal4*) transgene to drive a nuclear localized mCherry (*UAS-mCherry-nls*) reporter. *T2A-Gal4* lines carry insertions in genes such that *Gal4* expression is under the control of endogenous promoters and is driven in a pattern that most often recapitulates the expression of the gene in which the transgene is inserted^36, 37^. We costained *CG42458-T2A-Gal4>UAS-mCherry-nls* larval eye discs with Sens antibody, a known R8 marker (Fig. 3F-F’’’) and observe that mCherry is present in only one nucleus per ommatidium and colocalizes with Sens. Although mCherry expression starts few columns later than *sens* expression, these data show that *CG42458* is a novel R8-specific marker, and that the FeaturePlot accurately represents the endogenous expression pattern of *CG42458*. Similarly, *asense* (*ase*), *CG42313* and *sidestep* (*side*) FeaturePlots show expression in the same cluster as *CG42458*, and are likely novel R8 markers. To our knowledge, none of these four genes have been previously reported to be expressed in R8 cells. We also identified many novel markers for all other cell types and validated several *in vivo* (Fig. 3G, Supplementary Figs. 2-4 and Supplementary Table 1).

**Fig. 3:**
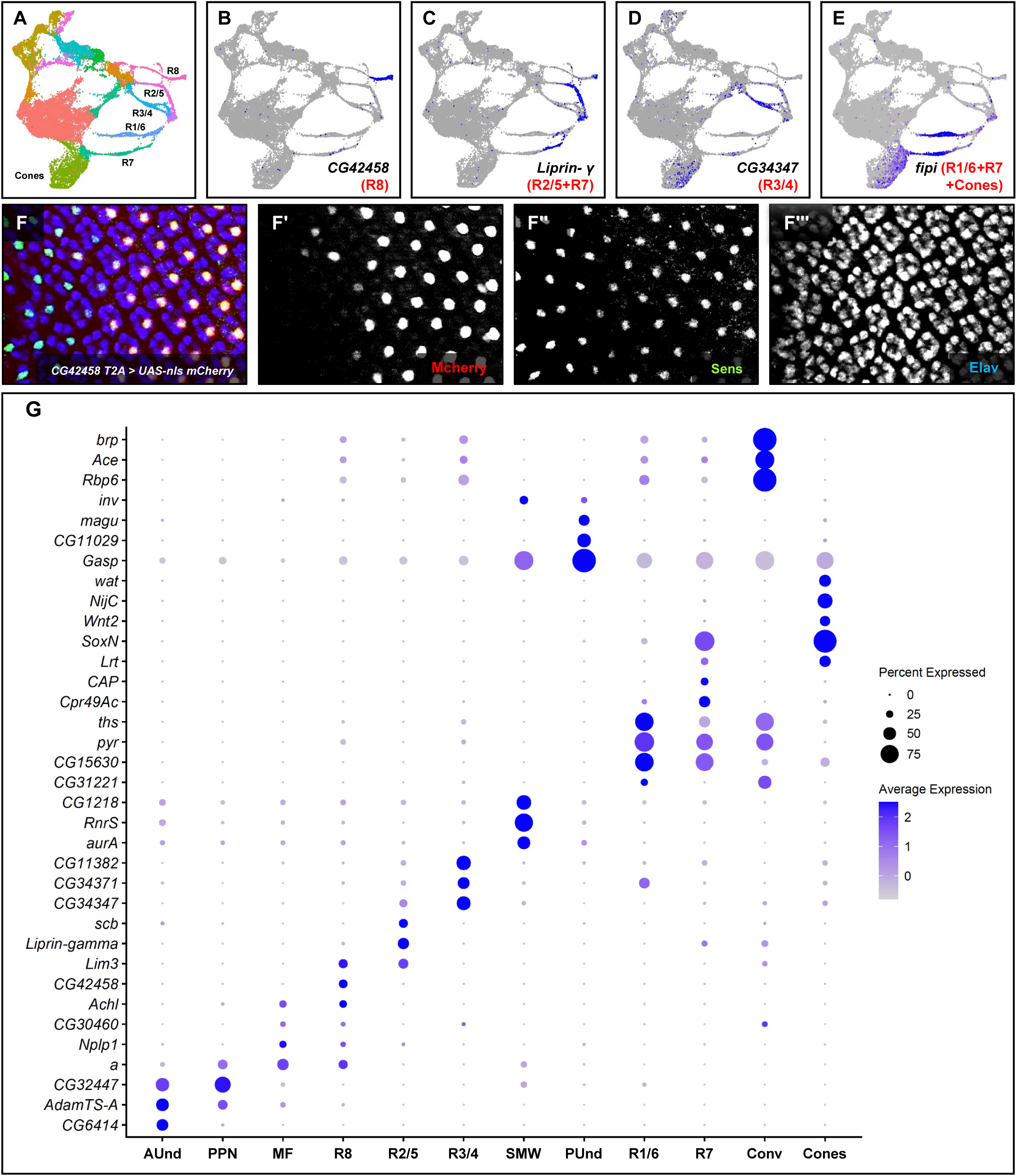
Identification and *in vivo* validation of novel markers. **A.** UMAP cluster plot of the late larval eye disc. **B-E.** FeaturePlots showing the expression of novel cell type-specific genes. **B.** *CG42458* FeaturePlot showing expression in R8. **C.** *liprin-γ* mRNA is detected in R2/5 and R7. **D.** *CG34347* is predominantly expressed in the R3/4 stream. Some expression is also detected in R2/5 and cones. **E.** *fipi* expression is observed in R1/6, R7 and a substantial fraction of cone cells. **F-F’’’.** Staining of eye discs from *CG42458-T2A-Gal4*>*UAS-nls-mCherry* larvae showing mCherry expression in Sens-positive cells. Note that *CG42458* is not expressed in early R8 cells (the left side of Panel F’). **G.** DotPlot of newly identified markers that are highly specific for each cell type present in the late larval eye disc.

**Fig. 4:**
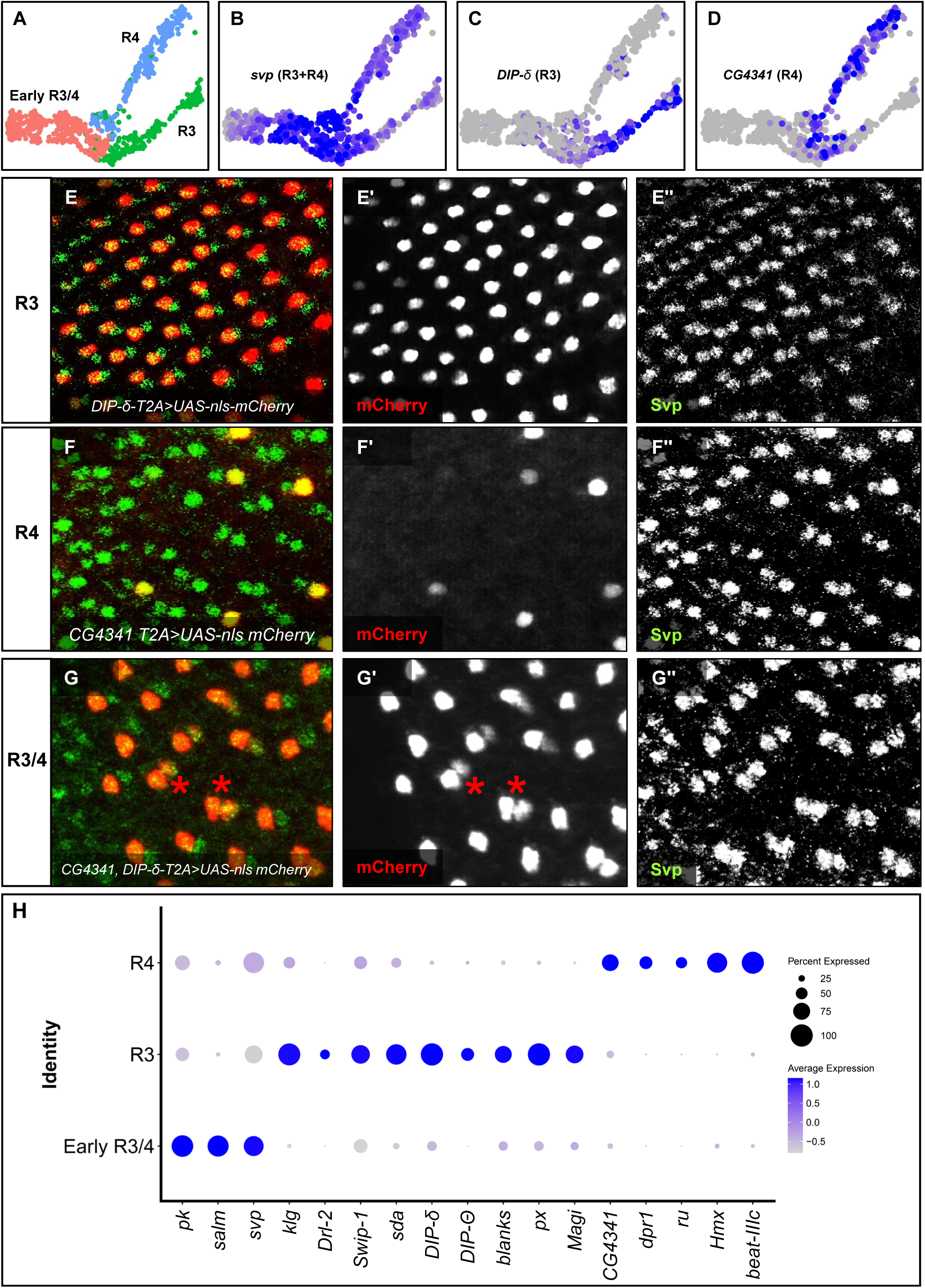
Novel markers that distinguish the R3 and R4 photoreceptors. **A.** UMAP cluster plot showing a subset of the R3/4 photoreceptor stream. The R3/4 stream splits into two distinct subclusters. **B.** FeaturePlot of *svp* showing expression in both the R3 and R4 streams after the split. **C.** *DIP-δ* FeaturePlot showing R3-specific expression. **D.** *CG4341* expression is highly specific to the R4 cluster. **E-E’’.** Larval eye disc image of *DIP-δ-T2A-Gal4>UAS-nls mCherry* stained with mCherry (red) and Svp (green). Only the anterior (left) of the two Svp-positive cells in each ommatidium costains with mCherry. **F-F’’.** *CG4341-T2A Gal4>UAS-nls-mCherry* eye discs stained with mCherry (red) and Svp (green). mCherry is also detected in one cell per ommatidium (although not in all ommatidia), but this costains with the posterior (right) of the two Svp-positive cells in an ommatidium. **G-G’’**. Larval eye disc carrying both *CG4341* and *DIP-δ-T2A-Gal4* cassettes costained with mCherry (red) and Svp (green). A pair of cells show mCherry expression (asterisk in **G** and **G’**) which also costain with Svp. **H.** DotPlot showing markers expressed early in the R3/4 stream as well as R3- and R4-specific novel markers.

### Novel markers that distinguish the R3 and R4 Photoreceptors

Our scRNA UMAP cluster plot shows that the R3/4 cluster emerges from the MF cluster as a single stream but then splits into two smaller streams (Fig. 1D, red arrow). We hypothesized that the R3/4 stream may be splitting into distinct R3 and R4 subtype clusters. Since the two sub-streams are apparent in both biological replicates, it seemed possible that the split is not an artifact of dimension reduction. We sub-clustered the R3/4 stream and generated a UMAP plot that shows three major clusters (Fig. 4A). These clusters resemble the R3/4 stream shown in Fig. 1D with a single cluster splitting into two sub-clusters. Differential gene expression analyses identified several genes that are specifically expressed in only one of the two split sub-clusters.

The genes *prickled* (*pk*), *svp* and *salm* are expressed in both R3 and R4^30, 38, 39^ a few columns posterior to the MF (Fig. 2F,J and Fig. 4B,H). We termed the cluster expressing these genes as ’Early R3/4.’ Notably, one of the split sub-clusters shows higher *E(spl)* gene expression (Supplementary Fig. 5B-E). Since Notch signaling is known to be higher in R4 than R3, we hypothesize this sub-cluster to be R4, while the other could represent R3 cells (Fig. 4A). Differential gene expression analyses identified several novel R3- or R4-specific markers (Fig. 4H), including *Dpr-interacting protein δ* (*DIP-δ*) (R3) and *CG4341* (R4).

In larval eye discs, R3 and R4 cells are immediately adjacent, with R4 being more posterior than R3 in the posterior part of the disc (Fig. 1B’)^40^. We drove *UAS-mCherry-nls* using *DIP-δ-T2A- Gal4* and costained larval eye discs with mCherry and Svp. If *DIP-δ* is an R3-specific marker, then we would expect mCherry and Svp coexpression in one cell per ommatidium that is anterior to the other Svp-expressing cell and this is precisely what our staining results show (Fig. 4E-E’’). These results suggest that *DIP-δ* is a novel R3-specific marker (Fig. 4E-E’’).

We obtained similar results when we stained *CG4341-T2A-Gal4* driven *UAS-mCherry-nls* eye discs (Fig. 4F-F’’). However, in contrast to *DIP-δ*, the mCherry and Svp coexpressing cell is posterior to the other Svp expressing cell, suggesting that it is an R4 cell and that *CG4341* is a novel R4 marker (Fig. 4F-F’’). We also drove *UAS-mCherry-nls* in larval eye discs carrying both *DIP-δ-* and *CG4341-T2A-Gal4* transgenes and observe mCherry and Svp in two cells per ommatidium (Fig. 4G-G’’). These data provide strong evidence that these genes are indeed novel R3- and R4-specific markers. Based on their FeaturePlot expression patterns, the genes shown in Fig. 4H are likely to be additional novel R3- or R4-specific markers. Taken together, these data demonstrate that we successfully profiled gene expression at very high resolution for all cell types present in the larval eye disc.

### Photoreceptor cells cluster as temporal streams converging on a common transcriptome

Our data show that each PR cluster appears as a thin stream emerging from the MF (R8, R2/5, R3/4) or PUnd cells (R1/6, R7) and all converge at the most mature (posterior/right) ends of the streams to a single transcriptional identity (Figs. 1D and 5A). Although the R7 and R8 streams do not directly contact the Convergence cluster, their gene expression patterns at their far right ends (i.e., most mature) largely mimic those seen in the Convergence cluster. To investigate the temporal pattern of PR development with higher resolution, we sub-clustered the MF and PRs and generated a UMAP cluster plot that shows the five streams corresponding to R8, R2/5, R3/4, R1/6 and R7 as well as the MF and Convergence clusters (Fig. 5A). The R8, R2/5 and R3/4 streams all emerge from the MF cluster, while the R1/6 and R7 stream origins are distinct as they are derived from undifferentiated cells following the SMW. Moreover, all five PR streams appear to merge into a common transcriptional identity on the posterior (right) side of the cluster plot (i.e., the Convergence cluster).

**Fig. 5:**
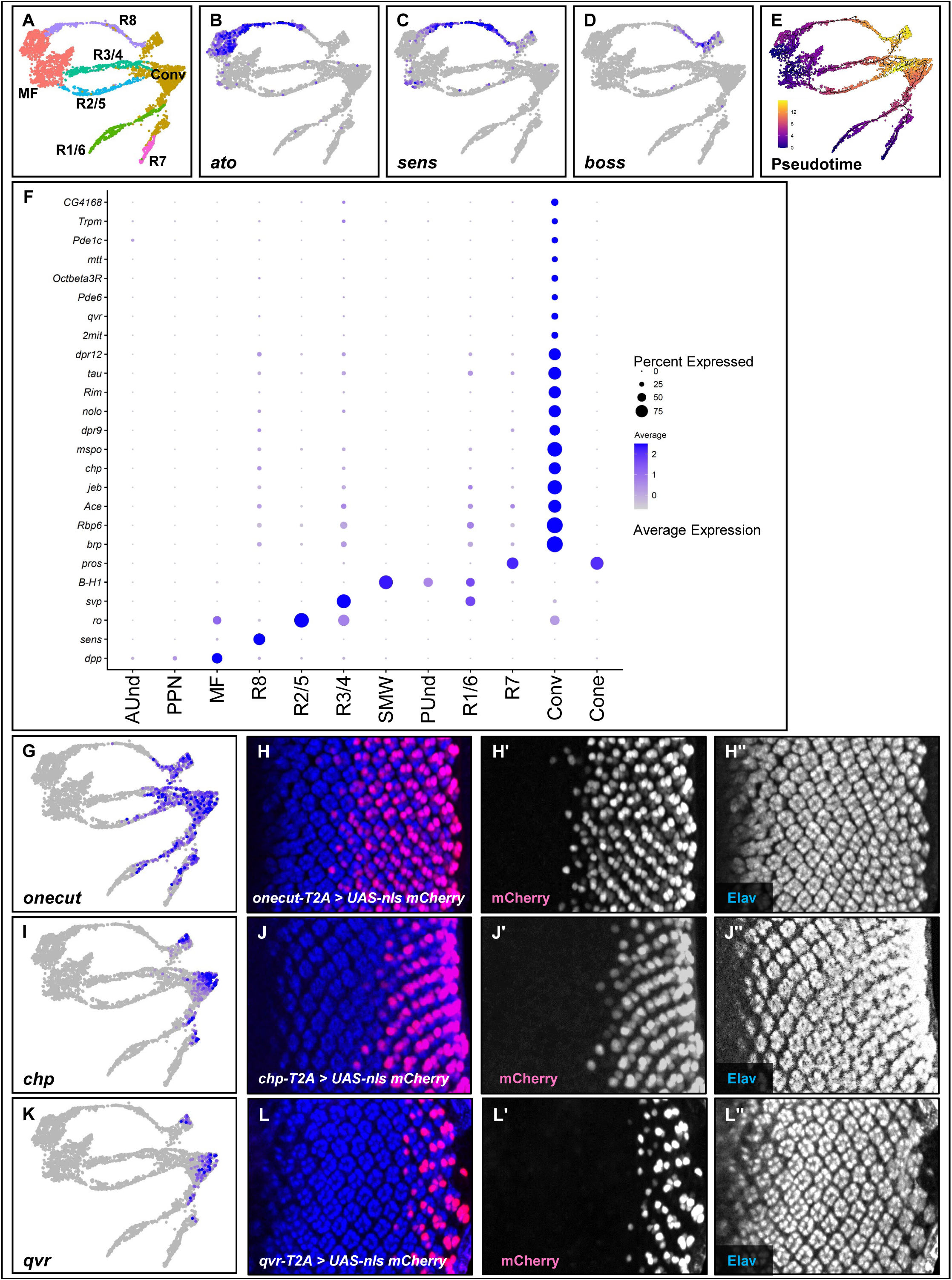
Distinct streams of photoreceptor cells emerge from undifferentiated clusters and then converge upon a common transcriptome. **A.** UMAP cluster plot showing only the MF and PRs. The PR clusters exhibit a spatiotemporal dimension with less mature PRs near the MF and more mature PRs near the Convergence cluster. **B.** FeaturePlot showing the expression of *ato* in the MF and R8 near the MF. C. FeaturePlot showing *sens* expression distributed throughout most of the R8 stream. **D.** FeaturePlot showing *boss* expression in more mature R8 PRs. **E.** Trajectory analysis of the MF and PRs showing early to late pseudotime along each PR stream. Purple indicates early pseudotime and late pseudotime is denoted as yellow. **F.** DotPlot showing expression of PR-, cone- and Convergence-specific markers. In the Convergence cluster, PRs have less distinct individual transcriptional identities with greatly reduced expression of subtype-specific markers. **G.** FeaturePlot showing *onecut*, which is expressed in late PRs and the Convergence cell cluster. **H-H’’**. *onecut-T2A-Gal4*>*UAS-nls-mCherry* larval eye disc costained with mCherry (red) and Elav (blue). mCherry expression begins six to seven columns of ommatidia posterior to the MF. **I.** FeaturePlot showing expression of *chaoptin* (*chp*) specifically in the Convergence cluster; no expression is observed in late PR streams. **J-J’’.** Colabeling of *chp-T2A-Gal4*>*UAS-mCherry- nls* with mCherry (red) and Elav (blue). mCherry is detected in a few posterior columns of ommatidia, consistent with the *chp* FeaturePlot. **K.** FeaturePlot of *quiver* (*qvr*) shows expression in the posterior (right) half of the Convergence cluster. **L-L’’**. *qvr-T2A-Gal4>UAS-nls-mCherry* eye disc costained with mCherry (red) and Elav (blue). mCherry is detected in just four ommatidial columns at the posterior margin of the eye disc.

We first compared the FeaturePlot expression patterns of the R8 markers *ato*, *sens,* and *boss* (Fig. 5B-D) with their *in vivo* expression patterns^25–27^. We observe a clear temporal progression of developmental identity along the R8 stream from anterior to posterior (left to right). Specifically, *ato*-expressing cells are within and immediately adjacent to the MF cluster, while *boss*-expressing cells are located at the opposite (posterior/right) end of the stream; *sens* expression is flanked by *ato* and *boss* along the stream. These FeaturePlot patterns strongly correlate with the known *in vivo* expression profiles of each gene, suggesting that the streams represent developmental trajectories of individual PRs with cells arranged as a function of time. Developmentally older PRs are located at the posterior ends (i.e., the right side in Figs. 1D and 5A) of each stream while younger cells (i.e., early PRs) are anterior (i.e., left), near the MF. Consistent with this interpretation, we find markers that exhibit the same spatiotemporal dimension in other PR streams as well. For example, *svp* is expressed in R1/6 in the first few columns posterior to the MF and is absent from R1/6 PRs located more posteriorly^30^. *B-H1* expression in R1/6 expression starts at about the same developmental time (i.e., distance from the MF) that *svp* expression stops in R1/6^41^. The FeaturePlots of *svp* and *B-H1* (Supplementary Fig. 6B,C) accurately represent these *in vivo* expression patterns with *B-H1* RNA detected at the posterior/right side of the R1/6 stream while *svp* RNA is in the anterior/left portion of the stream, with only minimal overlap between the two (black arrows, Supplementary Fig. 6B,C). We also performed trajectory analyses and observe a clear trend of early to late pseudotime along each PR stream (Fig. 5E). These data collectively show that the PR streams are two-dimensional developmental trajectories of individual PRs with cells computationally ordered from early to late along each stream.

Our UMAP cluster plots also show that most PR streams merge into a single Convergence cluster as they mature (at the posterior/right end of each stream) (Figs. 1D and 5A). Moreover, most PR subtype-specific gene expression (e.g., *sens*, *ro* and *svp*) strongly decreases in the Convergence cluster, PRs appear to lose at least some of their individual transcriptional identities, and instead express many markers related to axonogenesis, axon-pathfinding and synapse formation (Fig. 5F). Differential gene expression analyses identified 1,987 Convergence cluster marker genes (Supplementary Table 1). Among these, 69 are Class 1 genes that show strong expression specifically in the Convergence cluster (Fig. 5F and Supplementary Table 1). Since our scRNA data shows mature cells on the right of the UMAP plot, we expect converging PRs to be present in the posterior-most columns of the eye disc. For example, FeaturePlots of *onecut*, *chaoptin* (*chp*) and *quiver* (*qvr*), all show expression in the far right of the cluster plot, including the Convergence cluster (Fig. 5G,I,K). We tested if these expression patterns are observed *in vivo* by driving *UAS-mCherry-nls* using either *onecut*-, *chp*- or *qvr-T2A-Gal4* constructs and costained eye discs with mCherry and Elav (a pan-neuronal marker). Consistent with the FeaturePlots, we observe mCherry only in posterior PR columns and this costains with Elav for all three *T2A-Gal4* lines (Fig. 5H-H’’, 5J-J’’ and 5L-L’). Moreover, expression of these genes is progressively restricted from the posterior/right end of the PR streams to the distal tip of the Convergence cluster. Specifically, *onecut* is expressed in late PR streams and the Convergence cluster, *chp* is expressed only in the Convergence cluster but not in PR streams, and *qvr* is expressed only on the posterior/right side of the Convergence cluster. Again, the *in vivo* mCherry patterns resemble the expression patterns observed in FeaturePlots, with progressive restriction of expression to more posterior columns of the eye disc (Fig. 5G,I,K). These data suggest that our FeaturePlots are an accurate two-dimensional representation of the *in vivo* expression patterns of genes in the Convergence cluster (and distal tips of the R7 and R8 streams).

### A single nuclear chromatin accessibility atlas of the developing *Drosophila* eye

To complement our scRNA-seq dataset, we performed single nuclear assay for transposase accessible chromatin using sequencing (snATAC-seq) of late larval eye discs using the 10x Chromium Single Cell ATAC Reagent kit (Fig. 6A). We generated sequence data from three biological replicates and the raw data from all three were combined and analyzed, yielding about 29,000 cells. We then used Signac^42^ to perform QC, normalization, and dimension reduction steps. After removing non-retinal and peripodial cells, 20,035 cells remained, which is twice the number of cells in a single eye disc. A UMAP cluster plot was generated from these cells that shows all the expected cell identities present in the late larval eye disc (Fig. 6B).

**Fig. 6:**
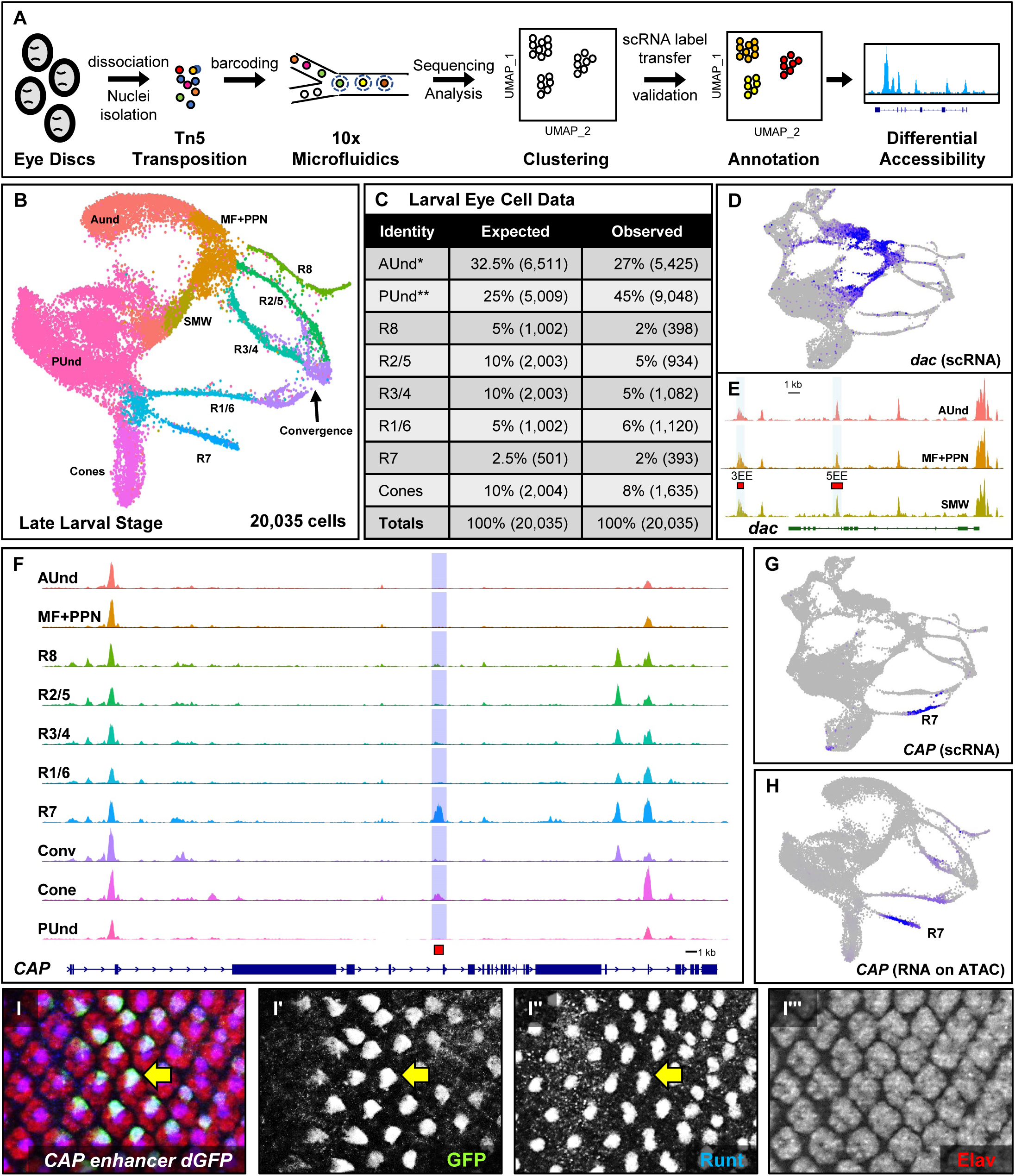
snATAC sequencing of late larval eye discs recapitulates the cellular identities observed in scRNA-seq. **A.** Schematic of snATAC-seq data generation and downstream analyses. **B.** UMAP cluster plot generated from snATAC sequencing of late larval eye discs shows all expected cell identities. Clusters of cells appear in a temporal progression similar to the UMAP plot from scRNA-seq. **C.** The expected and observed cell numbers and percentages for each cell type are shown. Peripodial cells were excluded and percentages were calculated for major cell types in the eye disc. **D.** scRNA-seq FeaturePlot showing *dac* mRNA expression in the AUnd, PPN, MF and SMW cell clusters. **E.** snATAC-seq CoveragePlot of the *dac* locus for the AUnd, MF+PPN and SMW clusters. The 3EE and 5EE *dac* enhancers (red bars) overlap snATAC-seq peaks in these clusters where *dac* is expressed. **F.** CoveragePlot of the *CAP* gene reveals a strong snATAC-seq peak highly specific for the R7 and, to a lesser degree, cone cell clusters. The red bar indicates the DNA fragment used to make the enhancer-reporter construct shown in **I-I’’’**. **G.** FeaturePlot of *CAP* mRNA showing R7-specific expression. **H.** FeaturePlot of *CAP* mRNA imputed on the snATAC-seq UMAP cluster plot showing mRNA distribution predominantly in R7. **I-I’’’.** Costaining of a larval eye disc from an animal carrying the *CAP* R7-specific peak DNA driving a destabilized GFP reporter with GFP (green), Run (blue), and Elav (red) antibodies. One cell in each ommatidium costains with GFP, Run, and Elav (yellow arrow, **I-I’’**).

Remarkably, the arrangement of clusters in the snATAC-seq UMAP plot (Fig. 6B) is highly similar to the clusters seen in the scRNA-seq UMAP plot (Fig. 1D). Like scRNA-seq, the snATAC-seq PR clusters appear as streams of cells, closely resembling the physical eye disc. Overall, the two dimensional arrangement of all clusters in the snATAC-seq data is nearly identical to that observed from scRNA-seq, including differentiating PRs, anterior and posterior undifferentiated cells, as well as the MF, SMW, PPN, and cone cell clusters. PRs appear as thin streams of cells that emerge from the MF or PUnd clusters and the R1-R6 streams converge into a common Convergence cluster. Taken together, these snATAC-seq data represent all of the expected cell types with good representation of each (Fig. 6C).

### snATAC-seq cluster identification and validation

Annotation and analysis of snATAC-seq data presents several challenges compared to scRNA-seq data and the limitations of the current snATAC technology make capturing the entire accessibility profile from an individual cell difficult^43^. snATAC-seq data are relatively sparse and may not reveal cellular variability at many individual regulatory elements, making cluster annotation challenging. We therefore used several approaches to classify and validate cell identities in our snATAC-seq cluster plots. First, we performed integrative analyses to classify snATAC-seq clusters based on cluster information from scRNA-seq data derived from eye discs at the same developmental stage (Fig. 1D). Using this strategy, cell identity labels were transferred from scRNA-seq to the snATAC-seq cluster map. This method has been used to map cell identities in snATAC-seq clusters from scRNA-seq data in *Drosophila*, humans, and mice^19, 44, 45^. In addition to transferring labels, gene expression values from scRNA-seq can also be transferred and imputed on snATAC-seq clusters such that marker gene expression patterns can be visualized on snATAC-seq cluster plots. Based on known gene expression patterns, we were able to validate predicted labels on the snATAC-seq clusters by coembedding scRNA-seq and snATAC-seq datasets to visualize both on the same UMAP plot^46^ (Supplementary Fig. 7).

To test the validity of our snATAC-seq data, we used previously published eye-specific enhancers to see if our data shows accessible chromatin corresponding to these enhancers. The *dachshund* (*dac*) gene is expressed in the MF, PPN and SMW (Fig. 2J and Fig. 6D). A 3’ enhancer (named ’3EE’) and a 5’ enhancer (named ’5EE’) recapitulate the endogenous *dac* pattern of expression in eye discs^47^. The *dac* snATAC-seq CoveragePlot shows an intronic peak and a 3’ peak, which correspond precisely with the 5EE and 3EE enhancers (red bars, Fig. 6E and Supplementary Fig. 8A,A’). Further, the 3’ peak is predominantly accessible in the AUnd, MF, and SMW clusters, reflecting the endogenous pattern of *dac* expression. Similarly, our snATAC-seq dataset shows accessible peaks that correspond to known eye enhancers for *ato*, *sens*, *lozenge*, *shaven* (*sv*), and *pros*^48–52^ (Fig. 7A,B,B’ and Supplementary Fig. 8).

**Fig. 7:**
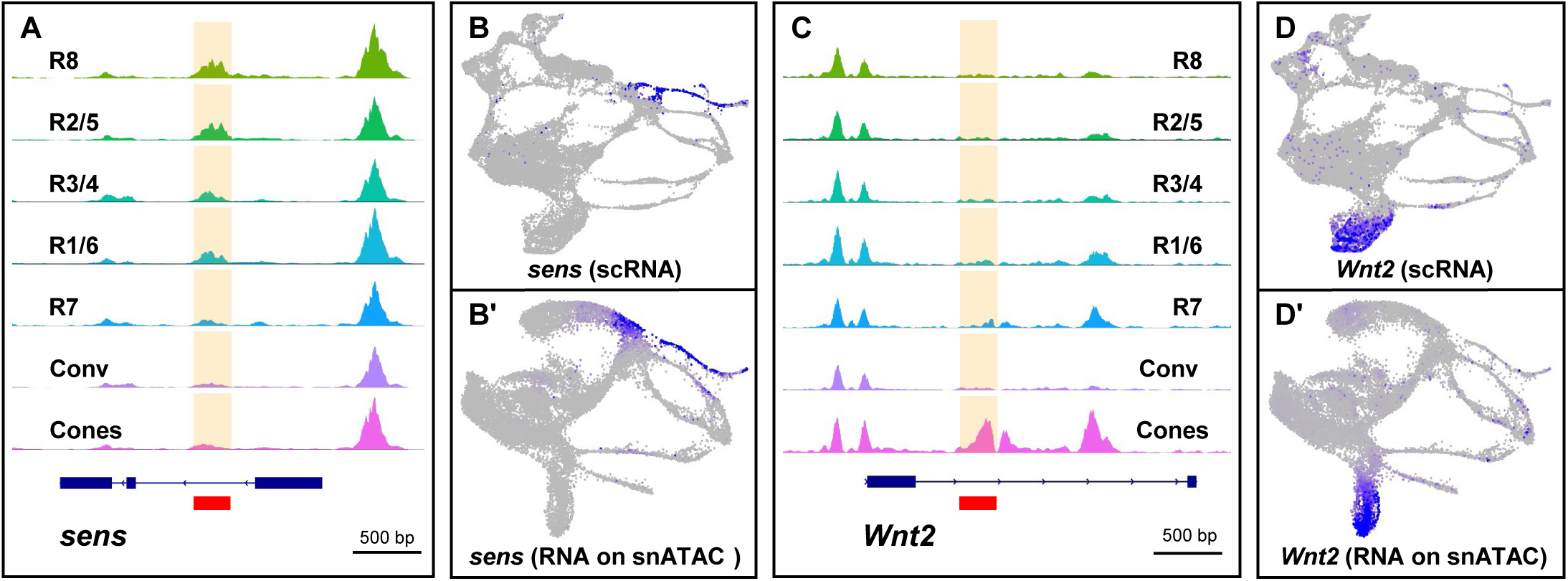
Chromatin is more accessible in photoreceptors compared to cones. **A.** snATAC-seq CoveragePlot showing the *sens* locus. The solid red bar denotes the *sens* F2 enhancer that is both necessary and sufficient for R8-specific expression *in vivo*. The snATAC-seq peak is highlighted in yellow. The s*ens F2* region is accessible in other PRs in addition to R8. **B.** scRNA-seq FeaturePlot of *sens* showing specific expression in the MF and R8. **B’.** *sens* mRNA imputed on a snATAC-seq UMAP plot showing distribution of *sens* mRNA specifically in the MF and R8. **C.** *Wnt2* CoveragePlot showing a cone-specific peak (solid red bar). The peak is accessible only in the cone cell cluster. **D.** scRNA-seq FeaturePlot of *Wnt2* showing mRNA expression is specific to the cone cell cluster. **D’.** *Wnt2* RNA overlay on the snATAC-seq cluster plot shows *Wnt2* expression is specific to cones.

We also used the JASPAR database^53^ to find overrepresented DNA motifs in snATAC-seq peaks. As one example, our motif analyses show overrepresentation of the *shaven* (*sv*) motif in both the R7 and cone cell clusters (Supplementary Fig. 9A-D). *sv* is an R7- and cone-specific marker which regulates neural and cone cell fate decisions in the eye disc^48^. Taken together, these observations suggest that our snATAC-seq data is of high quality, correspond well with published data, and that many of the cell type assignments have been validated *in vivo*.

To test if our snATAC-seq data predicts functional enhancer elements, we selected several cell type-specific peaks and used the corresponding DNA to drive reporter gene expression *in vivo*. The *CAP* gene scRNA-seq and snATAC-seq FeaturePlots show expression specifically in the R7 stream (Fig. 6G-H). Differential accessibility tests of the R7 cell cluster identify a peak in the fifth intron of the *CAP* gene as one of the top peaks. Moreover, the *CAP* locus genomic track reveals a peak that is specific and accessible primarily in the R7 cluster (Fig. 6F). To test if the DNA encompassing this peak contains an enhancer that is sufficient to drive reporter expression specifically in R7, we made reporter constructs with *destabilized GFP* (*dGFP*)^54^ driven by this peak DNA and costained transgenic larval eye discs with GFP, Run and Elav antibodies. We observe dGFP expression in a single cell per ommatidium that begins a few columns posterior to Elav expression and costains with the apical Run-positive R7 cell (Fig. 6I- I’’’). (58). These results show that the fifth intron snATAC-seq peak of *CAP* uncovers a novel R7- specific enhancer. Similarly, *2mit* mRNA is confined specifically to the Convergence cluster (Supplementary Fig. 9F,G) and differential accessibility region analyses of the snATAC-seq Convergence cluster reveals a peak in the third intron of the *2mit* gene. The CoveragePlot of *2mit* shows that this region is more accessible in the Convergence cluster compared to other cell identities (Supplementary Fig. 9E). We generated transgenic flies carrying *2mit* snATAC-seq peak DNA upstream of *dGFP*, and stained larval eye discs with GFP and Elav. As expected, the peak-region DNA carries a Convergence cluster-specific enhancer. We observe dGFP expression only in the most posterior columns in late larval eye discs, and dGFP colocalizes with Elav-positive cells (Supplementary Fig. 9H-H’’). Taken together, these data show that our snATAC-seq data can accurately predict *in vivo* enhancer activity and can be used to identify novel enhancers with temporal and cell-specific properties. Furthermore, these data also validate the cell identity assignments made via label transfer from scRNA-seq to snATAC-seq datasets. We next performed trajectory analyses of the snATAC-seq dataset by choosing AUnd cells as the root cells. Although the the pseudotime graphs of snATAC-seq (Supplementary Fig. 9I) and scRNA-seq (Fig. 1E) are very similar, one striking difference is apparent: PRs that differentiate prior to the SMW (R8, R2/5 and R3/4) in the snATAC-seq data show a much earlier profile compared to those in the scRNA-seq data. This may be a reflection of the generally more permissive chromatin observed for the PRs compared to cells that differentiate following the SMW (R1/6, R7, and cones). These results suggest that the chromatin accessibility dynamics of R1/6, R7 and cones may be different from other PR subtypes.

### Chromatin accessibility differences between PRs and non-neuronal cone cells

To explore chromatin accessibility across snATAC-seq cell clusters, we performed differential accessibility tests between cell clusters using default metrics to generate lists of differentially accessible marker peaks for each cell cluster in our snATAC-seq data set (Supplementary Table 2). We observe that the number of differentially accessible peaks is higher in PR cell clusters than undifferentiated cells with the AUnd, MF and PUnd cell clusters showing 2-4 fold fewer differentially accessible peaks than PRs. We also investigated the chromatin profile of PRs and cone cells to determine if there are accessibility differences between neuronal and non-neuronal differentiating cells. We analyzed the top 100 differentially accessible peaks in PR clusters and found that only 15% (or fewer) of peaks are subtype-specific (Supplementary Figs. 10 and 11); instead, most of the top 100 peaks are present in nearly all PRs. We also examined accessibility profiles near several known PR markers (Fig. 2A-I) and found a similarly permissive chromatin profile across all PRs with little or no specificity for the subtype in which the marker is expressed. For instance, the *sens* F2 enhancer is both necessary and sufficient for R8-specific expression of *sens*. However, the CoveragePlot of *sens* shows that the F2 enhancer region is open in most PRs and not just R8 cells (red box, Fig. 7A). These data suggest that PR differentiation may be regulated by subtype-specific transcription factor activity and that chromatin may be in a largely permissive state with only 3% of PR genes appearing to show specific snATAC-seq peaks (Supplementary Table 2).

In contrast, among the top 100 snATAC-seq peaks in the cone cell cluster, 70% are predominantly specific to the cone cell identity. For instance, *Wnt2* mRNA is detected only in the cone cell cluster (Fig. 7D,D’) and a snATAC-seq peak in the sole intron of *Wnt2* (red bar, Fig. 7C) is one of the top 100 differentially accessible peaks. Moreover, the peak is accessible only in the cone cell cluster. These results suggest that chromatin may be more restrictive in the cone cell cluster and cone cell differentiation may involve both localized TF activity as well as restrictive chromatin. Taken together, these data suggest that there may be functionally important chromatin accessibility differences between PRs and non-neuronal cone cells in the larval eye disc.

### Single Cell regulatory Network Inference and Clustering (SCENIC) analyses of scRNA-seq data reveals putative novel cell type-specific regulators

We used the SCENIC tool to identify important regulators and gene regulatory networks from our scRNA-seq data^55^. SCENIC identifies coexpression modules (termed ’regulons’) that comprise sets of genes coexpressed with transcription factors. Our SCENIC results show that PRs cluster separately from other cell types (Fig. 8), suggesting that PRs share similar regulatory networks and cell states that are distinct from other cell types in the eye disc.

**Fig. 8:**
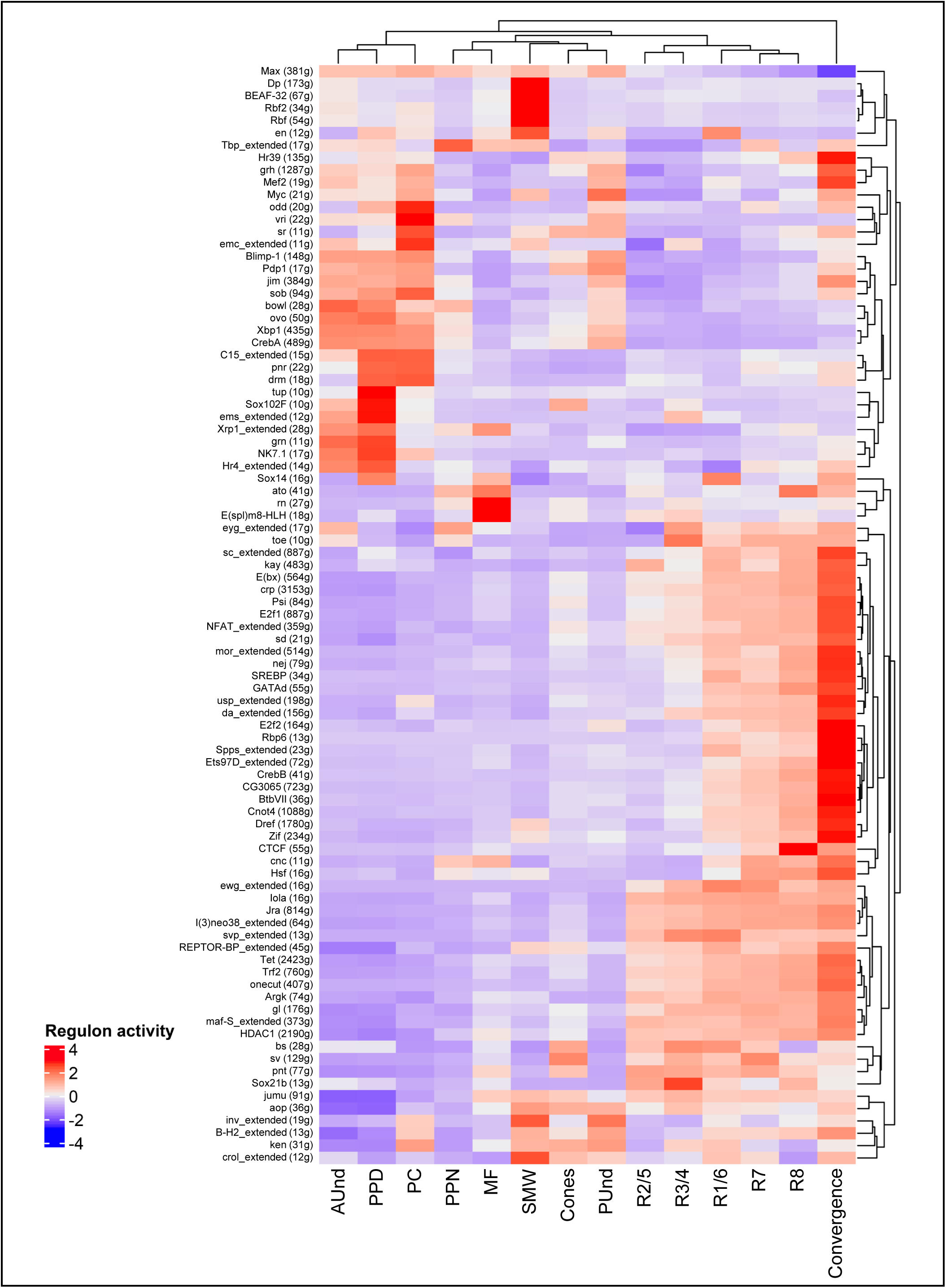
SCENIC analyses reveals known and novel putative regulators of each cell identity. Heatmap showing top regulators of each cell cluster in scRNA-seq. The intensity of red denotes the activity of regulons. The Convergence cell cluster is distinct with a large number of upregulated regulons compared to all other cell identities.

Furthermore, the observed regulons recapitulate many of the known spatiotemporal gene regulatory network dynamics observed during larval eye disc differentiation. For instance, we observe that the *ato* and *E(spl)m8-HLH* regulons are most active in the MF^28, 56^ and *svp* in R3/4 and R1/6^30^. We also identified top putative regulators for each cluster by calculating the regulon specificity score (RSS), which is based on the entropy, expression level, and specificity of each regulon in a given cluster (Supplementary Fig. 12A). The regulons *CTCF*, *ato*, *erect wing* (*ewg*), and *RNA binding protein 6* (*Rbp6*) are among the top R8 regulons. Both *ato* and *ewg* are well known R8 regulators^28, 57^, but *CTCF* and *Rbp6* are novel putative R8 regulators. Similarly, the regulons *kayak* (*kay*), *longitudinals lacking* (*lola*), *Jun-related antigen* (*Jra*) and *ewg* are among the top R2/5 regulators, whereas *svp*, *ewg* and *lola* are top R3/4 regulators. *kay*, *lola* and *Jra* have been implicated in eye development^58–60^, and *svp* and *lola* are known to regulate R3/4 fate choice^30, 60^. The top putative regulators for all cell clusters are shown in Supplementary Fig. 12A. Interestingly, the Convergence cell cluster on the SCENIC heatmap comprises a remarkably large number of regulons highly upregulated in this cluster compared to all other cell identities (Fig. 8). Our analyses identified *Rbp6, ewg* and *onecut* as top Convergence cluster regulators and expression of *Rbp6* is highly specific to the Convergence cell cluster and late R1/6 cells (Supplementary Fig. 12B). Our snATAC-seq motif analyses using the JASPAR and i-cisTarget databases^53, 61^ also show Onecut motif enrichment in the Convergence cluster. *onecut* is highly conserved and required for horizontal cell development in mice^62–64^. In *Drosophila*, we have found that *onecut* null mutants show age-related retinal dysmorphology with loss of PR rhabdomeres (to be reported elsewhere). Using PANTHER^65^, we performed Gene Ontology (GO) term enrichment analyses with the genes included in each Convergence cell cluster regulon and found that GO terms related to axon guidance (GO:0007411) and neuron projection guidance (GO:0097485) are highly enriched. In summary, SCENIC analyses identified many known regulators of eye development and thereby provide further validation of our scRNA-seq data. In addition, several potentially important novel regulators were also identified for each cell cluster, which provide new avenues to investigate the regulatory mechanisms underlying eye development.

## Discussion

In this study, we report a comprehensive and high quality single cell genomics atlas of the developing *Drosophila* late larval eye disc with more than two-fold coverage of the number of cells present in a single disc. Our dissection and dissociation steps were performed in the presence of Actinomycin D (ActD), which inhibits transcription, thereby minimizing stress-related changes to transcription and chromatin configuration^66–68^. Therefore, our scRNA-seq and snATAC-seq data most likely reflect the endogenous gene expression and chromatin configuration of all known cell types in the larval eye disc. Our scRNA-seq data was derived from 26,999 high quality and viable cells from eye discs with a sequencing depth of 2.1 billion reads, while our snATAC-seq data was generated from 20,595 high quality nuclei with a sequencing depth of 680 million reads. Our data show distinct cell clusters corresponding to all major cell identities present in the eye disc and each cell type is well represented (Fig. 1D and Fig. 7C). Furthermore, both our scRNA-seq and snATAC-seq UMAP cluster plots show distinct cell clusters that are ordered sequentially and in a manner consistent with their developmental age. In particular, differentiating photoreceptor cell clusters emerge from the MF (R8, R2/5 and R3/4) or PUnd (R1/6, R7 and cones) clusters as distinct streams. Moreover, as inferred from known marker gene expression (Fig. 5), each PR stream represents a developmental continuum with cells arranged progressively along the stream with newly differentiating PRs near the MF and more mature PRs at the far right (posterior) of each stream. Remarkably, this arrangement of PRs and other clusters closely resembles the position of cell types in the physical eye disc. Therefore, our data sets comprise a computational representation of the larval eye disc such that the FeaturePlot can be used as a virtual *in situ* to reveal the expression patterns of most genes in the eye disc without the need for *in vivo* staining or spatial mapping software. The expression and distribution of known subtype-specific markers and snATAC-seq peaks that correspond to known eye-specific enhancers in our data further validate this point. It is also of note that cluster plots generated from two completely different types of data (scRNA-seq and snATAC-seq) portray nearly identical arrangements of cell clusters and both are highly reminiscent of the cell type distributions in physical eye disc. In addition, analyses of our data reveal many putative novel markers, enhancers and regulators for each cell type, particularly for PR subtypes and cones, several of which have been validated *in vivo*. Further, precursor cells of different PRs and cones cluster separately and show distinct transcriptomic and chromatin accessibility profiles. Investigating the profiles of undifferentiated cell clusters and functional testing of genes in precursor cell clusters may reveal signaling mechanisms and pathways that underlie the specification and differentiation of distinct cell types in the eye disc and are likely to benefit research groups that use the larval eye disc as a model system.

Unexpectedly, we do not observe obvious dorsal-ventral (DV) clustering in our datasets. Specifically, although the gene *mirror* (*mirr*) is known to be highly expressed in the dorsal half of late larval eye discs^69^, we do not observe distinct clustering of *mirr*-expressing cells in our data and instead observe *mirr* expression throughout all clusters (Supplementary Fig. 1C). One possibility is that our cellular coverage and sequencing depth may be insufficient to detect transcriptional variation between dorsal and ventral cells. Alternatively, although some genes are clearly expressed either dorsally or ventrally, there are too few such genes at this stage of development to drive clustering of cells with specific dorsal or ventral identities.

Shortly after R3 and R4 begin to differentiate, dorsal and ventral ommatidia rotate 90° in opposite directions and appear as mirror images of each other in adults. Establishment of this ommatidial orientation requires Fz/Dsh and N signaling between R3 and R4^16, 17^. The polarizing signal is initiated in R3 through the transcriptional activation of the N ligand Delta (Dl) by Fz/Dsh. As a result, N is activated in the neighboring R4 cell and therefore has higher N signaling compared to R3. The *Enhancer of split* (*E*(*spl*)) genes are the downstream effectors of N signaling and a ∼500 bp enhancer fragment (named mδ0.5) of *E(spl)mδ-HLH* is the only marker known to differentiate R3 and R4 and drives reporter expression in an R4-specific pattern^16^. Our scRNA-seq cluster plot shows that the R3/4 PR stream splits into two smaller streams, which we have identified as R3 and R4. As expected, our data shows higher expression of *E*(*spl*) genes in R4 compared to R3. We also identified several novel R3- and R4- specific markers and validated *DIP-δ* and *CG4341* as R3- and R4-specific, respectively.

Interestingly, these markers are detected in the smaller streams but are not detected in the early R3/4 cluster. We confirmed these expression patterns *in vivo* for both *DIP-δ* and *CG4341*: reporter gene expression is observed in R3 or R4 in posterior ommatidial columns, but not in early R3/4 cells. This suggests that R3 and R4 are transcriptionally very similar in the first few columns posterior to the MF, but become distinct and express different markers further posteriorly. In contrast, our snATAC-seq clustering does not show any R3/4 distinction with the R3/4 stream appearing as a single stream without a split. This suggests that the chromatin profiles of the R3 and R4 streams may be very similar. To the best of our knowledge, this is the first report that identifies genes that distinguish the R3 and R4 photoreceptors and these may provide insights into mechanisms that establish epithelial polarity. We have not yet identified any genes that distinguish individual PRs in the R2/5 and R1/6 PR pairs in our analyses. However, since individual members of these PR pairs are not known to possess distinct functions (unlike R3 and R4), the lack of a transcriptional distinction within these PR pairs may not be unexpected.

Our scRNA-seq and snATAC-seq clustering results show that PRs R1-6 (and, perhaps to a lesser extent, R7 and R8) have reduced subtype identity as they mature, converge at the posterior regions of the eye disc, and express markers related to axon projection, guidance and synapse formation. These PRs therefore exhibit higher transcriptional diversity during early stages of differentiation and then become more similar in the Convergence cluster. Since the Convergence cluster appears in independent data sets that measure very different aspects of the genome (scRNA-seq and snATAC-seq), it seems unlikely that the Convergence cluster is an artifact of dimension reduction. Moreover, like PR streams, cells in the Convergence cluster also appear to be ordered in a temporal series that strongly correlates with *in vivo* gene expression. We have also identified many markers and putative regulators that are expressed in the Convergence cluster, including *Rbp6* and *onecut*, which are well-conserved among animals and play important roles in neural development^70, 71^. The phenomenon of neurons losing their identity and converging on a common transcriptome during synaptogenesis was reported for *Drosophila* olfactory projection neurons and the optic lobe^13, 72^. The biological significance of why neurons undergo convergence is not clear, but it has been hypothesized that synapse formation in the brain beyond neuropil targeting may require synchronous wiring of neurons and therefore all such neurons must converge and express similar markers^72^. A similar requirement may pertain to PRs that are projecting their axons to the medulla and lamina of the optic lobe. Early in development, PR transcriptional distinction may be required for precise positioning of each PR within a single ommatidium. As PRs mature, however, synchronous targeting may be necessary for proper synaptogenesis and therefore may demand highly similar transcriptomes.

Furthermore, since R1-6 and R7/8 target their axons to different layers of the brain, it may be expected that R7 and R8 do not fully converge compared to R1-6. Future functional studies using genes and regulators identified in the Convergence cluster reported in this study may help unravel the mechanisms and biological significance of converging neurons. The convergence phenomenon may be a common theme in *Drosophila* neural development and, to our knowledge, this is the first such report for larval photoreceptors.

Our snATAC-seq data shows that there are considerable differences in the chromatin accessibility of PRs and cones. While cones show restrictive chromatin with several snATAC-seq peaks accessible only in the cone cell cluster, the chromatin in PRs appears to be more permissive, with most peaks accessible in most or all PR subtypes, including several peaks that are associated with known marker genes that show very specific subtype expression (e.g. *sens- F2*). Furthermore, cone cell cluster snATAC-seq peaks are often near genes that show cone-specific expression, which is distinct when compared to PR subtype clusters, which collectively show peaks near genes involved in axon guidance and projection (Supplementary Table 3).

Since the chromatin states of PR subtypes are very similar, cell type differentiation may be largely controlled by distinct transcription factor activity in different PR subtypes. In contrast, both the chromatin state and localized cone-specific gene expression may be involved in cone differentiation. We also observe that the snATAC-seq peaks associated with PRs are generally not accessible in cone cells, further highlighting the differences in cell states between these two major cell types. We also observe that restrictive chromatin increases in cells that differentiate after the SMW with cones showing the most specific peaks, followed by R7 and R1/6, suggesting that chromatin is substantially remodeled after cells exit the SMW. Although the mechanisms underlying differences in the chromatin states of PRs and cones is unclear, these results highlight the potential utility of our snATAC-seq data in deciphering cell fate determination and differentiation.

In summary, we provide a high quality and extensive single cell genomics atlas of the *Drosophila* larval eye that represents all cell types in the eye disc, including identification of many novel cell type-specific genes, enhancers and putative regulators. Moreover, photoreceptor clusters are observed as streams representing developmental continuums that strongly correlate with *in vivo* gene expression. Intriguingly, these initially distinct PR streams converge upon a common transcriptomic identity by late larval stages. Our analyses of chromatin profiles of photoreceptors and cones suggest differences that may underlie distinct mechanisms of differentiation of neuronal and non-neuronal cells. These single cell resources provide a wealth of genome-wide transcriptomic and chromatin accessibility data that will dramatically enable novel approaches to investigating mechanisms of cell fate determination, development, and function.

## Methods

### Fly husbandry

All flies used in this report were maintained at 25°C on cornmeal agar medium. We obtained the following fly stocks from the Bloomington *Drosophila* Stock Center: *UAS-mCherry-nls* (38424), *MiMIC-CG42458* (67472), *CRIMIC-Liprin-gamma* (79357), *MiMIC-CG34347* (76674), *MiMIC-DIP- δ* (90320), *MiMIC-CG4341* (76620), *CRIMIC-chp* (78931), and *CRIMIC-qvr* (86367). The *roFF::GFP* flies were generated by *GenetiVision Corporation* with support from grant R44 GM148146 and based on the protocol described in Manivannan et al, 2019^73^.

### Dissociation of late larval eye discs into single cells for scRNA-seq

We collected 25 to 30 eye discs from *Canton-S* late larvae (0 hrs after puparium formation) and immediately transferred them to a LoBind 1.5 ml Eppendorf tube containing 700 µl ice-cold Rinaldini solution supplemented with 1.9 µM Actinomycin-D, a known transcription inhibitor.

After dissection, 16 µl of collagenase (100 mg/ml; Sigma-Aldrich #C9697) and 2 µl of dispase (1mg/ml; Sigma-Aldrich #D4818) were added to the tube. The tube was then placed horizontally in a shaker and the eye discs were dissociated for 50 min at 32°C at 250 rpm. The solution was pipetted every 10 min to disrupt clumps of cells. The cells were then diluted with 1 ml of Rinaldini solution containing 0.05% Bovine Serum Albumin (BSA). The diluted cell suspension was passed through a 35 µm sterile filter and centrifuged at 4°C at 50 to 100 rcf to obtain a cell pellet. The pellet was washed once with Rinaldini + 0.05% BSA and subjected to centrifugation. The cell pellet was resuspended in Rinaldini + 0.05% BSA and the viability was assessed using Hoechst-propidium iodide solution. Samples that showed >95% viability were used for scRNA-seq experiments.

### Single cell RNA-seq using 10x Genomics

We used Chromium Next GEM Single Cell 3’ Reagent Kit v3.1 from 10x Genomics to generate single cell libraries from single cell suspensions that showed >95% viability and at 1000-1200 cells/µl concentration. Briefly, single cell suspensions were loaded on a 10x Genomics Chromium Controller along with Gel Beads containing barcoded primers and oil emulsion. The Chromium Controller isolates each cell in an oil droplet with a Gel Bead (GEM). The cells are lysed and the mRNA is captured and barcoded within the oil-bead emulsion followed by reverse transcription to synthesize cDNA. The cDNAs from each cell were pooled and a library was generated and sequenced with a NovaSeq 6000 (Illumina). We sequenced two biological replicates to a sequencing depth of ∼1 billion reads each. FASTQ files generated from each sequencing run were combined and analyzed using the Cell Ranger v6.0.1 aggr (aggregate) pipeline. The *Drosophila melanogaster* reference genome Release 6 (dm6) was used to make the reference genome using the Cell Ranger ’mkref’ pipeline.

### Seurat Analyses

The filtered gene expression matrices from the Cell Ranger output had ∼36,000 cells with a sequencing depth of 2.6 billion reads. These cells were used as input to perform downstream analyses in Seurat v4.03. We first removed potential multiplets and lysed cells by retaining cells that showed a total number of genes between 200 and 5000 and cells that showed low mitochondrial gene percentage (<30%). The filtered cells were then processed with the Seurat SCTransform algorithm that normalizes and scales the data across all cells. The number of variable features used for SCTransform was 5000 and regression was performed using mitochondrial genes to remove effects of mitochondrial gene expression on clustering. The dimensionality of the data was then reduced using the top 50 dimensions, and the data was clustered using the RunUMAP, FindNeighbors and FindClusters functions in Seurat. Using known markers, we removed cells from antenna (*Distal-less* (*dll*))^74^, glia (*reversed polarity* (*repo*))^75^ and brain (*found in neurons* (*fne*))^76^, and 26,999 cells with a sequencing depth of ∼2.1 billion reads were retained from the eye disc proper, Oc, PC and LM, and the PPD. Differential marker gene lists for all cell clusters were generated using the FindAllMarkers function with a log-fold change threshold value of 0.25 and a minimum percentage of cells in which the gene is detected of 25%. For trajectory analyses, ’SeuratWrappers’ R package was used to convert the scRNA-seq Seurat object to a celldataset object for analyses in Monocle 3. The cells were ordered in pseudotime by selecting the AUnd cell cluster as the root cells. Similar analyses were performed for the MF and PR subclusters with the MF as the root cells.

### Dissociation of larval eye discs into single nuclei for snATAC-seq

We dissected and collected 30 to 40 late larval eye discs from *Canton-S* and removed antennal discs prior to transferring them to a LoBind Eppendorf tube containing ice-cold 1x PBS supplemented with 1.9 µM ActD. The 1x PBS was replaced with 100 µl lysis buffer containing digitonin (0.005%) and Nonidet P40 Substitute (Sigma, #74385) and incubated on ice for 5 min. The solution was pipetted every 1 min to lyse the cell membranes and to release intact nuclei into the solution. After 5 min, the lysis reaction was stopped by diluting with Tris-HCl (10 mM, pH 7.4) wash buffer containing 10% BSA. The solution was then passed through a 10 µm filter to remove cell debris and clumps and subjected to centrifugation at 4°C. The supernatant was discarded and the nuclear pellet was resuspended in Tris-HCl wash buffer and centrifuged one more time at 4°C to obtain a nuclear pellet. The pellet was then resuspended in 1x Nuclei Buffer (10x Genomics, PN-2000153, PN-2000207) and the integrity of nuclear membranes was ascertained using bright field microscopy. Nuclear samples that showed intact nuclear membranes without blebbing were used for snATAC-seq experiments.

### snATAC-seq using 10x Genomics and downstream analyses

We used 10x Chromium Next GEM Single Cell ATAC Kit v2 to generate libraries from dissociated nuclei. Briefly, nuclei at a concentration of 3,000/µl were used to recover ∼10,000 cells for each snATAC-seq experiment. The nuclear suspension was first mixed with the 10x Chromium Transposition Mix and incubated at 37°C for 30 min for the transposition reaction. The Transposase Mix fragments the DNA and adds adapter sequences to the fragments. The transposed nuclei were then loaded into the 10x Chromium Controller along with barcoded Gel Beads and partitioning oil to generate GEMs. The Gel Beads were then dissolved and the barcoded DNA fragments pooled to generate libraries. We generated libraries from three independent biological replicates and sequenced them separately with a NovaSeq 6000 (Illumina). FASTQ files generated from each sequencing run were pooled and analyzed using the cellranger-atac aggr v2.0 pipeline. The output of the Cell Ranger ATAC pipeline was used as input for Signac v1.8.0 for quality control (QC) and downstream analyses. We first removed potential multiplets and lysed nuclei by selecting nuclei that showed total peak region fragments between 500 and 30,000. Next, we selected nuclei that showed more than 25% (15% default in Signac) of fragments in peaks, which represents the fraction of all fragments that fall within ATAC-seq peaks. Reads that may be artifactual (blacklist regions) were also removed using the ENCODE dm6 blacklist^77^. The filtered data was then subjected to normalization and dimensional reduction followed by the identification of clusters using the RunUMAP function. We used the top 50 dimensions to perform dimension reduction. Cell clusters were annotated by integrating with scRNA-seq and transferring cell labels from scRNA-seq UMAP plots to snATAC-seq clusters. This procedure involved computing gene activities from snATAC-seq followed by identification of transfer anchors between scRNA-seq and snATAC-seq data by canonical correlational analysis. Cell identity annotations were then transferred from scRNA-seq to snATAC-seq clusters using integration anchors. In addition, gene expression values from scRNA-seq data were transferred and imputed on snATAC-seq clusters and marker gene expression patterns were visualized on the snATAC-seq cluster plot. Cell clusters pertaining to the brain, glia, PC and PPD were removed and only cells from the eye disc proper were retained.

### Immunohistochemistry

Late larval eye discs were dissected and immediately transferred to a 1.5 µl Eppendorf tube containing ice-cold 1x PBS and fixed in 3.7% paraformaldehyde in PBS for 30 min at room temperature. Eye discs were then washed 3 times with PBS+0.3% Triton X-100 (PBT) and blocked using 5% normal goat serum in PBT. Primary antibody incubations were done overnight at 4°C. Secondary antibody incubations were performed at room temperature for at least 1 hr.

Eye discs were washed and mounted on glass slides for imaging. A Zeiss Apotome Imager microscope was used to generate optically stacked images, which were processed with Zen Blue and Adobe Photoshop software. We used the following antibodies: rat anti-Elav (RRID:AB, #52818, 1:500), chicken anti-GFP (RRID:AB, #300798, 1:1000), rabbit anti-mCherry (RRID:AB, #2889995, 1:2000), guinea pig anti-Runt (gift from Dr. Claude Desplan), guinea pig anti-Sens (a gift from Hugo Bellen, 1:1000) and mouse anti-Svp (RRID:AB, #2618079, 1:500). The following secondary antibodies were used at 1:500 concentration: Cy5 anti-rat (RRID: AB, #2534067), Cy5 anti-guinea pig (RRID:AB, #2340460), Alexa 488 anti-guinea pig (RRID: AB, #2534117), Alexa 488 anti-chicken (RRID:AB, #2762843), Alexa 568 anti-rabbit (RRID:AB, #2534017), Alexa 488 anti-mouse (RRID: AB, #2536161) and Alexa 555 anti-rat (RRID: AB, #2535855).

### Transgenic assays to identify functional enhancers

To identify functional enhancers, we selected cell type-specific peaks and designed primers that span the entire ’called’ peak sequences. We PCR amplified the DNA corresponding to the peaks and cloned the purified PCR products into *pH-Stinger-dGFP-attB* or *pH-Stinger-mCherry-attB* vectors. We generated transgenic flies using site-specific integration and used the attP2 landing site. Transgenic flies were generated by *GenetiVision Corporation*. Transgenic late larval eye discs were dissected and stained with the antibodies listed in the results as previously described^78^. Eye discs were imaged using a Zeiss Apotome Imager microscope to generate optically stacked images. Zen blue and Adobe Photoshop software were used to process the stacked images.

### GO term analyses

Genes that were near differentially accessible PR, Convergence and cone cluster peaks were used for analysis with Panther. The fold enrichment of the top enriched GO terms for biological processes were used to make bar graphs.

## Data availability

All raw and processed data are being uploaded onto Gene Expression Omnibus, Accession number: XXXX). Seurat .rds files are available upon request.

## Code availability

All R scripts used to generate the data shown in this work will be uploaded onto GitHub

## Acknowledgements

We thank Claude Desplan for sharing anti-Runt antibodies and Hugo Bellen for sharing anti- Senseless antibodies. We thank Nick Tran, Shinya Yammamoto, Justin Ma and Benjamin Frankfort for their insightful comments on our manuscript. This work was funded in part by The Retina Research Foundation. Library prep and sequencing was performed at the Single Cell Genomics Core at Baylor College of Medicine and was partially supported by NIH shared instrument grants (S10OD023469, S10OD025240), P30EY002520 and CPRIT grant RP200504.

## Author Contributions

K.K.B.R. and K.Y. performed the larval eye disc dissection and analyses of scRNA-seq data.

The single-cell dissociation protocol and snATAC-seq analyses were performed by K.K.B.R.

Y.L. performed cDNA library construction and sequencing of single-cell cDNA libraries.

K.K.B.R. prepared and conducted the immunofluorescence imaging of larval eye discs.

Y.S. performed enhancer cloning.

The manuscript was prepared by K.K.B.R. and reviewed by K.Y., R.C., and G.M.

## Correspondence

Correspondence should be sent to Graeme Mardon.

## References

1. Karaiskos, N. et al. The Drosophila embryo at single-cell transcriptome resolution. Science 358, 194–199 (2017).

2. Consortium, T. M. Single-cell transcriptomics of 20 mouse organs creates a Tabula Muris. Nature 562, 367–372 (2018).

3. He, S. et al. Single-cell transcriptome profiling of an adult human cell atlas of 15 major organs. Genome biology 21, 1–34 (2020).

4. Travaglini, K. J. et al. A molecular cell atlas of the human lung from single-cell RNA sequencing. Nature 587, 619–625 (2020).

5. Yamagata, M., Yan, W. & Sanes, J. R. A cell atlas of the chick retina based on single-cell transcriptomics. Elife 10, e63907 (2021).

6. Jeibmann, A. & Paulus, W. Drosophila melanogaster as a model organism of brain diseases. International journal of molecular sciences 10, 407–440 (2009).

7. Prüßing, K., Voigt, A. & Schulz, J. B. Drosophila melanogaster as a model organism for Alzheimer’s disease. Molecular Neurodegeneration 8, 35, doi:10.1186/1750-1326-8-35 (2013).

8. Yamaguchi, M. & Yoshida, H. in Drosophila Models for Human Diseases (ed Masamitsu Yamaguchi) 1-10 (Springer Singapore, 2018).

9. Sang, T.-K. & Jackson, G. R. Drosophila models of neurodegenerative disease. NeuroRx 2, 438–446 (2005).

10. Hanson, I. & Van Heyningen, V. Pax6: more than meets the eye. Trends in Genetics 11, 268–272 (1995).

11. Wawersik, S. & Maas, R. L. Vertebrate eye development as modeled in Drosophila. Human Molecular Genetics 9, 917–925 (2000).

12. Donner, A. L. & Maas, R. L. Conservation and non-conservation of genetic pathways in eye specification. International Journal of Developmental Biology 48, 743–753 (2004).

13. Li, H. et al. Classifying Drosophila Olfactory Projection Neuron Subtypes by Single-Cell RNA Sequencing. Cell 171, 1206–1220.e1222, doi:10.1016/j.cell.2017.10.019 (2017).

14. Davie, K. et al. A single-cell transcriptome atlas of the aging Drosophila brain. Cell 174, 982–998. e920 (2018).

15. Yeung, K. et al. Single cell RNA sequencing of the adult Drosophila eye reveals distinct clusters and novel marker genes for all major cell types. Communications Biology 5, 1370, doi:10.1038/s42003-022-04337-1 (2022).

16. Cooper, M. T. & Bray, S. J. Frizzled regulation of Notch signalling polarizes cell fate in the Drosophila eye. Nature 397, 526–530 (1999).

17. Fanto, M. & Mlodzik, M. Asymmetric Notch activation specifies photoreceptors R3 and R4 and planar polarity in the Drosophila eye. Nature 397, 523–526 (1999).

18. Ariss, M. M., Islam, A. B. M. M. K., Critcher, M., Zappia, M. P. & Frolov, M. V. Single cell RNA- sequencing identifies a metabolic aspect of apoptosis in Rbf mutant. Nature Communications 9, 5024, doi:10.1038/s41467-018-07540-z (2018).

19. Bravo González-Blas, C., et al. Identification of genomic enhancers through spatial integration of single-cell transcriptomics and epigenomics. Molecular systems biology 16, e9438 (2020).

20. Wiegleb, G., Reinhardt, S., Dahl, A. & Posnien, N. Tissue dissociation for single-cell and single- nuclei RNA sequencing for low amounts of input material. Frontiers in Zoology 19, 27, doi:10.1186/s12983-022-00472-x (2022).

21. Brown, N. L., Sattler, C. A., Paddock, S. W. & Carroll, S. B. Hairy and emc negatively regulate morphogenetic furrow progression in the Drosophila eye. Cell 80, 879–887 (1995).

22. Seimiya, M. & Gehring, W. J. The Drosophila homeobox gene optix is capable of inducing ectopic eyes by an eyeless-independent mechanism. Development 127, 1879–1886 (2000).

23. Chanut, F. & Heberlein, U. Role of decapentaplegic in initiation and progression of the morphogenetic furrow in the developing Drosophila retina. Development 124, 559–567 (1997).

24. Pignoni, F. & Zipursky, S. L. Induction of Drosophila eye development by decapentaplegic. Development 124, 271–278 (1997).

25. Reinke, R. & Zipursky, S. L. Cell-cell interaction in the Drosophila retina: the bride of sevenless gene is required in photoreceptor cell R8 for R7 cell development. Cell 55, 321–330 (1988).

26. Baker, N. E., Yu, S. & Han, D. Evolution of proneural atonal expression during distinct regulatory phases in the developing Drosophila eye. Current Biology 6, 1290–1302 (1996).

27. Frankfort, B. J., Nolo, R., Zhang, Z., Bellen, H. & Mardon, G. senseless repression of rough is required for R8 photoreceptor differentiation in the developing Drosophila eye. Neuron 32, 403–414 (2001).

28. Jarman, A. P., Grell, E. H., Ackerman, L., Jan, L. Y. & Jan, Y. N. Atonal is the proneural gene for Drosophila photoreceptors. Nature 369, 398–400 (1994).

29. Kimmel, B. E., Heberlein, U. & Rubin, G. M. The homeo domain protein rough is expressed in a subset of cells in the developing Drosophila eye where it can specify photoreceptor cell subtype. Genes & Development 4, 712–727 (1990).

30. Mlodzik, M., Hiromi, Y., Weber, U., Goodman, C. S. & Rubin, G. M. The Drosophila seven-up gene, a member of the steroid receptor gene superfamily, controls photoreceptor cell fates. Cell 60, 211–224 (1990).

31. Higashijima, S.-i., et al. Dual Bar homeo box genes of Drosophila required in two photoreceptor cells, R1 and R6, and primary pigment cells for normal eye development. Genes & Development 6, 50–60 (1992).

32. Kauffmann, R. C., Li, S., Gallagher, P. A., Zhang, J. & Carthew, R. W. Ras1 signaling and transcriptional competence in the R7 cell of Drosophila. Genes & Development 10, 2167–2178 (1996).

33. Blochlinger, K., Jan, L. Y. & Jan, Y. N. Postembryonic patterns of expression of cut, a locus regulating sensory organ identity in Drosophila. Development 117, 441–450 (1993).

34. Daga, A., Karlovich, C. A., Dumstrei, K. & Banerjee, U. Patterning of cells in the Drosophila eye by Lozenge, which shares homologous domains with AML1. Genes Dev 10, 1194–1205, doi:10.1101/gad.10.10.1194 (1996).

35. Flores, G. V., Daga, A., Kalhor, H. R. & Banerjee, U. Lozenge is expressed in pluripotent precursor cells and patterns multiple cell types in the Drosophila eye through the control of cell- specific transcription factors. Development 125, 3681–3687 (1998).

36. Diao, F. et al. Plug-and-play genetic access to drosophila cell types using exchangeable exon cassettes. Cell Rep 10, 1410–1421, doi:10.1016/j.celrep.2015.01.059 (2015).

37. Lee, P.-T. et al. A gene-specific T2A-GAL4 library for Drosophila. Elife 7 (2018).

38. Rawls, A. S. & Wolff, T. Strabismus requires Flamingo and Prickle function to regulate tissue polarity in the Drosophila eye. Development 130, 1877–1887, doi:10.1242/dev.00411 (2003).

39. Domingos, P. M. et al. Regulation of R7 and R8 differentiation by the spalt genes. Developmental biology 273, 121–133 (2004).

40. Tomlinson, A. & Ready, D. F. Neuronal differentiation in the Drosophila ommatidium. Developmental Biology 120, 366–376, doi:https://doi.org/10.1016/0012-1606(87)90239-9 (1987).

41. Mavromatakis, Y. E. & Tomlinson, A. Switching cell fates in the developing Drosophila eye. Development 140, 4353–4361, doi:10.1242/dev.096925 (2013).

42. Stuart, T., Srivastava, A., Madad, S., Lareau, C. A. & Satija, R. Single-cell chromatin state analysis with Signac. Nature Methods 18, 1333–1341, doi:10.1038/s41592-021-01282-5 (2021).

43. Chen, H. et al. Assessment of computational methods for the analysis of single-cell ATAC-seq data. Genome Biology 20, 241, doi:10.1186/s13059-019-1854-5 (2019).

44. Pervolarakis, N. et al. Integrated single-cell transcriptomics and chromatin accessibility analysis reveals regulators of mammary epithelial cell identity. Cell reports 33, 108273 (2020).

45. Muto, Y. et al. Single cell transcriptional and chromatin accessibility profiling redefine cellular heterogeneity in the adult human kidney. Nature communications 12, 1–17 (2021).

46. Stuart, T. et al. Comprehensive integration of single-cell data. Cell 177, 1888–1902. e1821 (2019).

47. Pappu, K. S., et al. Dual regulation and redundant function of two eye-specific enhancers of the Drosophila retinal determination gene dachshund. (2005).

48. Fu, W., Duan, H., Frei, E. & Noll, M. shaven and sparkling are mutations in separate enhancers of the Drosophila Pax2 homolog. Development 125, 2943–2950, doi:10.1242/dev.125.15.2943 (1998).

49. Sun, Y., Jan, L. Y. & Jan, Y. N. Transcriptional regulation of atonal during development of the Drosophila peripheral nervous system. Development 125, 3731–3740 (1998).

50. Yan, H., Canon, J. & Banerjee, U. A transcriptional chain linking eye specification to terminal determination of cone cells in the Drosophila eye. Developmental Biology 263, 323–329, doi:https://doi.org/10.1016/j.ydbio.2003.08.003 (2003).

51. Hayashi, T., Xu, C. & Carthew, R. W. Cell-type-specific transcription of prospero is controlled by combinatorial signaling in the Drosophila eye. Development 135, 2787–2796, doi:10.1242/dev.006189 (2008).

52. Pepple, K. L., et al. Two-step selection of a single R8 photoreceptor: a bistable loop between senseless and rough locks in R8 fate. (2008).

53. Sandelin, A., Alkema, W., Engström, P., Wasserman, W. W. & Lenhard, B. JASPAR: an open- access database for eukaryotic transcription factor binding profiles. Nucleic acids research 32, D91–D94 (2004).

54. Atkins, M. et al. Dynamic rewiring of the Drosophila retinal determination network switches its function from selector to differentiation. PLoS Genet 9, e1003731, doi:10.1371/journal.pgen.1003731 (2013).

55. Aibar, S. et al. SCENIC: single-cell regulatory network inference and clustering. Nature Methods 14, 1083–1086, doi:10.1038/nmeth.4463 (2017).

56. Knust, E., Tietze, K. & Campos-Ortega, J. A. Molecular analysis of the neurogenic locus Enhancer of split of Drosophila melanogaster. The EMBO Journal 6, 4113–4123 (1987).

57. Hsiao, H.-Y., Jukam, D., Johnston, R. & Desplan, C. The neuronal transcription factor erect wing regulates specification and maintenance of Drosophila R8 photoreceptor subtypes. Developmental Biology 381, 482–490, doi:https://doi.org/10.1016/j.ydbio.2013.07.001 (2013).

58. Ciapponi, L., Jackson, D. B., Mlodzik, M. & Bohmann, D. Drosophila Fos mediates ERK and JNK signals via distinct phosphorylation sites. Genes Dev 15, 1540–1553, doi:10.1101/gad.886301 (2001).

59. Read, R. D., Bach, E. A. & Cagan, R. L. Drosophila C-terminal Src kinase negatively regulates organ growth and cell proliferation through inhibition of the Src, Jun N-terminal kinase, and STAT pathways. Mol Cell Biol 24, 6676–6689, doi:10.1128/mcb.24.15.6676-6689.2004 (2004).

60. Zheng, L. & Carthew, R. W. Lola regulates cell fate by antagonizing Notch induction in the Drosophila eye. Mechanisms of Development 125, 18–29, doi:https://doi.org/10.1016/j.mod.2007.10.007 (2008).

61. Herrmann, C., Van de Sande, B., Potier, D. & Aerts, S. i-cisTarget: an integrative genomics method for the prediction of regulatory features and cis-regulatory modules. Nucleic Acids Res 40, e114, doi:10.1093/nar/gks543 (2012).

62. Wu, F. et al. Onecut1 is essential for horizontal cell genesis and retinal integrity. J Neurosci 33, 13053–13065, 13065a, doi:10.1523/jneurosci.0116-13.2013 (2013).

63. Sapkota, D. et al. Onecut1 and Onecut2 redundantly regulate early retinal cell fates during development. Proceedings of the National Academy of Sciences 111, E4086–E4095 (2014).

64. Kreplova, M. et al. Dose-dependent regulation of horizontal cell fate by Onecut family of transcription factors. PLoS One 15, e0237403, doi:10.1371/journal.pone.0237403 (2020).

65. Mi, H. et al. The PANTHER database of protein families, subfamilies, functions and pathways. Nucleic acids research 33, D284–D288 (2005).

66. Perry, R. P. & Kelley, D. E. Inhibition of RNA synthesis by actinomycin D: characteristic dose- response of different RNA species. Journal of cellular physiology 76, 127–139 (1970).

67. Wadkins, R. M. & Jovin, T. M. Actinomycin D and 7-aminoactinomycin D binding to single- stranded DNA. Biochemistry 30, 9469–9478 (1991).

68. Wu, Y. E., Pan, L., Zuo, Y., Li, X. & Hong, W. Detecting activated cell populations using single- cell RNA-seq. Neuron 96, 313–329. e316 (2017).

69. McNeill, H., Yang, C.-H., Brodsky, M., Ungos, J. & Simon, M. A. mirror encodes a novel PBX- class homeoprotein that functions in the definition of the dorsal-ventral border in the Drosophila eye. Genes & development 11, 1073–1082 (1997).

70. Nguyen, D. N., Rohrbaugh, M. & Lai, Z. The Drosophila homolog of Onecut homeodomain proteins is a neural-specific transcriptional activator with a potential role in regulating neural differentiation. Mech Dev 97, 57–72, doi:10.1016/s0925-4773(00)00431-7 (2000).

71. Siddall, N. A. et al. Drosophila Rbp6 is an orthologue of vertebrate Msi-1 and Msi-2, but does not function redundantly with dMsi to regulate germline stem cell behaviour. PLoS One 7, e49810, doi:10.1371/journal.pone.0049810 (2012).

72. Özel, M. N. et al. Neuronal diversity and convergence in a visual system developmental atlas. Nature 589, 88–95, doi:10.1038/s41586-020-2879-3 (2021).

73. Manivannan, S. N., Pandey, P. & Nagarkar-Jaiswal, S. Flip-flop mediated conditional gene inactivation in Drosophila. Bio-protocol 9, e3157–e3157 (2019).

74. Dong, P. D., Chu, J. & Panganiban, G. Coexpression of the homeobox genes Distal-less and homothorax determines Drosophila antennal identity. Development 127, 209–216 (2000).

75. Xiong, W.-C., Okano, H., Patel, N. H., Blendy, J. A. & Montell, C. repo encodes a glial-specific homeo domain protein required in the Drosophila nervous system. Genes & development 8, 981–994 (1994).

76. Samson, M.-L. & Chalvet, F. found in neurons, a third member of the Drosophila elav gene family, encodes a neuronal protein and interacts with elav. Mechanisms of development 120, 373–383 (2003).

77. Amemiya, H. M., Kundaje, A. & Boyle, A. P. The ENCODE Blacklist: Identification of Problematic Regions of the Genome. Scientific Reports 9, 9354, doi:10.1038/s41598-019-45839-z (2019).

78. Au-Hsiao, H.-Y., et al. Dissection and Immunohistochemistry of Larval, Pupal and Adult Drosophila Retinas. JoVE, e4347, doi:doi:10.3791/4347 (2012).

