## Supplemental Figures 1-14 for "A Single Cell Genomics Atlas of the *Drosophila* Larval Eye Reveals Distinct Developmental Timelines and Novel Markers for All Photoreceptor Subtypes"

| **A. Sequencing Metrics** | |
|---|---|
| Total cell number | 26,999 |
| Means reads per cell | 76,014 |
| Number of reads | 2.1 billion |
| Median genes per cell | 2,173 |
| Fraction reads in cells | 94% |
| Median UMI counts per cell | 10,089 |

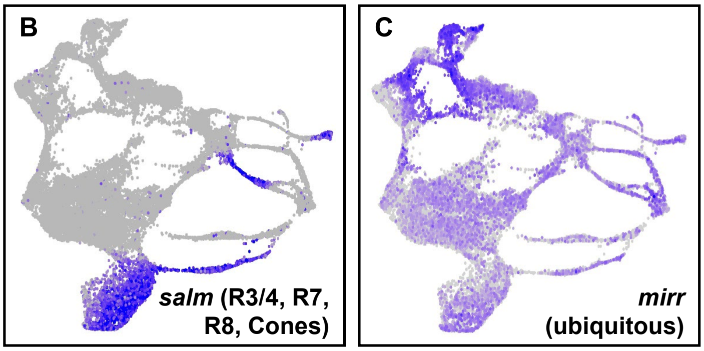


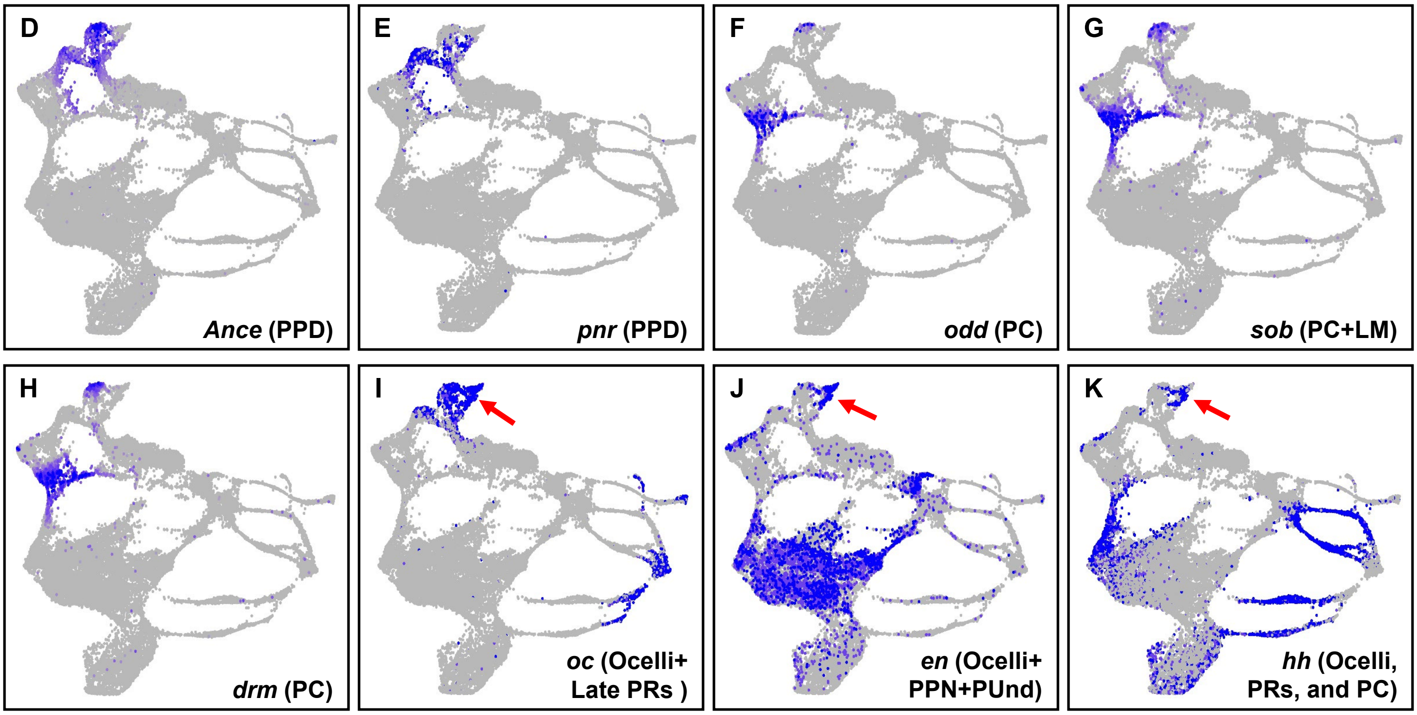

**Supplementary Fig. 1: scRNA-seq metrics and identification of the PPD and Oc cell clusters in scRNA-seq data. A.** Table showing quality metrics of the larval eye scRNA-seq data set. **B.** FeaturePlot showing *salm* expression in R3/4, R7, cones and late R8. The *salm* FeaturePlot expression is consistent with published studies. **C.** FeaturePlot of *mirr* showing ubiquitous expression without an apparent dorsal cluster. **D-K.** FeaturePlots of known marker genes used to identify the PPD, PC and OC clusters. **D.** FeaturePlot of the PPD marker *Angiotensin converting enzyme* (*Ance*) shows expression in the PPD. **E.** FeaturePlot for *pannier* (*pnr*) in the PPD cell cluster. **F-H.** FeaturePlots for *odd skipped* genes. *odd skipped* (*odd*, **F**), *sister of odd and bowl* (*sob*, **G**), and *drumstick* (*drm*, **H**) show expression specifically in the PC cell cluster. **I-K**. FeaturePlots showing expression of ocelli markers (red arrows): *ocelliless* (*oc*, **I**), *engrailed* (*en*, **J**), and *hedgehog* (*hh*, **K**).

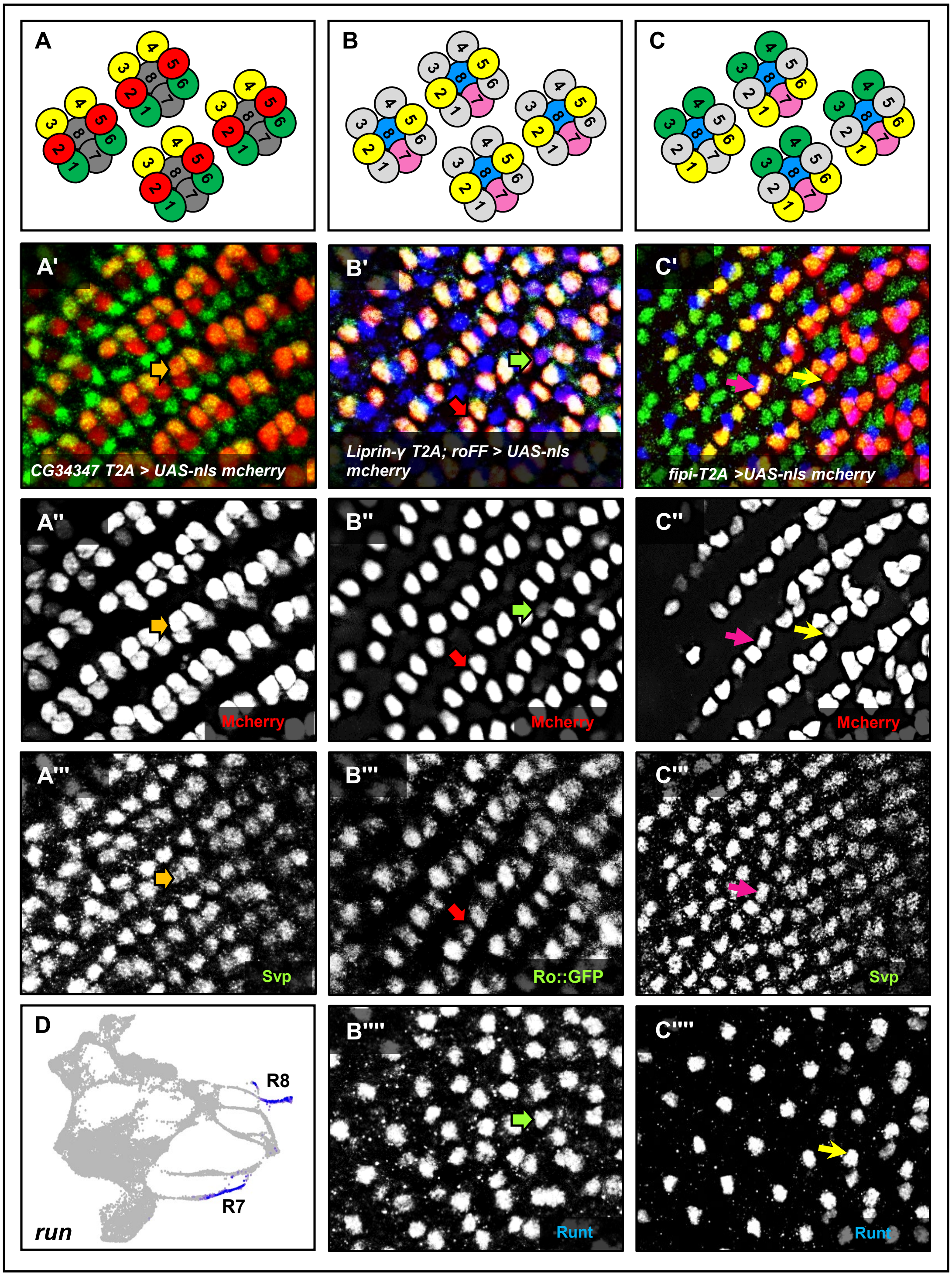
A
B
C
A'
CG34347 T2A > UAS-nls mcherry
B'
Liprin-γ T2A; roFF > UAS-nls mcherry
C'
fipi-T2A >UAS-nls mcherry
A''
Mcherry
B''
Mcherry
C''
Mcherry
A'''
Svp
B'''
Ro::GFP
C'''
Svp
D
R8
R7
run
B''''
Runt
C''''
Runt

**Supplementary Fig. 2: Identification and *in vivo* validation of novel markers.**

**A-C.** Schematics of four ommatidia with eight (R1-R8) photoreceptors each. The colors denote the expression observed in **A'-C'. A'-A'''.** *CG34347-T2A-Gal4>UAS-nls-mCherry* eye disc colabeled with mCherry (red) and Svp (green). Two Svp-positive cells also costain for mCherry (orange arrow in **A',A''** and **A'''**). **B'-B''''.** Colabeling of a *liprin-γ-T2A-Gal4; ro-flipflop-GFP>UAS-nls-mCherry* larval eye disc with mCherry (red), Run (blue) and GFP (green). *liprin-γ-T2A-Gal4* drives mCherry expression in R2/5 (red arrow in **B'-B'''**) and faintly in R7 (green arrow in **B'**, **B''** and **B'''**), consistent with the expression observed in the FeaturePlot (Figure 3C). **C'-C'''.** Costaining of a *fipi-T2A-Gal4>UAS-nls-mCherry* eye disc with mCherry (red), Svp (green) and Runt (blue). Two mCherry-positive cells costain with Svp (purple arrow in **C'-C'''**). A third cell costains with Run (yellow arrow in **C'**,**C''** and **C''''**). **D.** FeaturePlot for *run* shows expression in late R7 and R8.

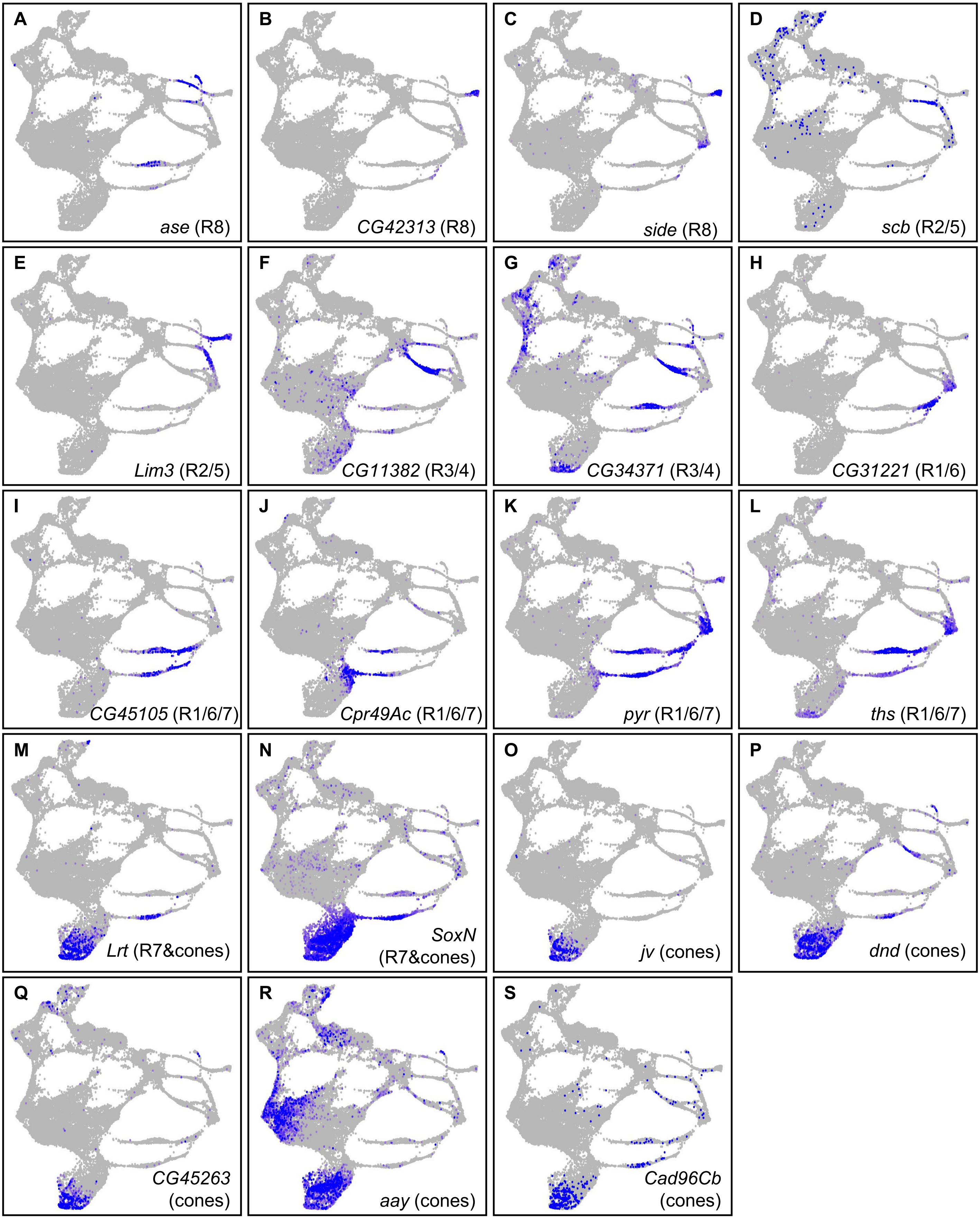
A
ase (R8)
B
CG42313 (R8)
C
side (R8)
D
scb (R2/5)
E
Lim3 (R2/5)
F
CG11382 (R3/4)
G
CG34371 (R3/4)
H
CG31221 (R1/6)
I
CG45105 (R1/6/7)
J
Cpr49Ac (R1/6/7)
K
pyr (R1/6/7)
L
ths (R1/6/7)
M
Lrt (R7&cones)
N
SoxN (R7&cones)
O
jv (cones)
P
dnd (cones)
Q
CG45263 (cones)
R
aay (cones)
S
Cad96Cb (cones)

**Supplementary Fig. 3: Identification of novel cell type-specific markers from scRNA-seq data. A-S.** FeaturePlots showing the expression and distribution of novel PR and cone cell cluster markers. *asense* (*ase*, **A**) and *CG42313* (**B**) are expressed in R8. **C.** *sidestep* (*side*) is predominantly expressed in late R8. *scab* (*scb*, **D**) and *Lim3* (**E**) show expression in the R2/5 cluster. *Lim3* is also expressed in R8. FeaturePlots of *CG11382* (**F**) and *CG34371* (**G**) show mRNA in R3/4. *CG34371* is also expressed in R1/6. **H.** FeaturePlot of *CG31221* showing expression in late R1/6 and the Convergence cluster. **I-L**. FeaturePlots of genes showing R1/6- and R7-specific expression: *CG45105* (**I**), *Cuticular protein 49Ac* (*Cpr49Ac*, **J**), *pyramus* (*pyr*, **K**), and *thisbe* (*ths*, **L**)*. pyr* and *ths* also show expression in the Convergence cluster. *Leucine-rich tendon-specific protein* (*Lrt*, **M**) and *SoxN* (**N**) are expressed in the R7 and cone cell clusters. **O-S.** FeaturePlots showing the expression of novel cone cell markers: *javelin* (*jv*, **O**), *dead end* (*dnd*, **P**), *CG45263* (**Q**), *astray* (*aay*, **R**), and *Cadherin 96Cb* (*Cad96Cb*, **S**). *aay* is also expressed in the PUnd cell cluster while *dnd* and *Cad96Cb* also show sporadic and weak expression in some PRs.

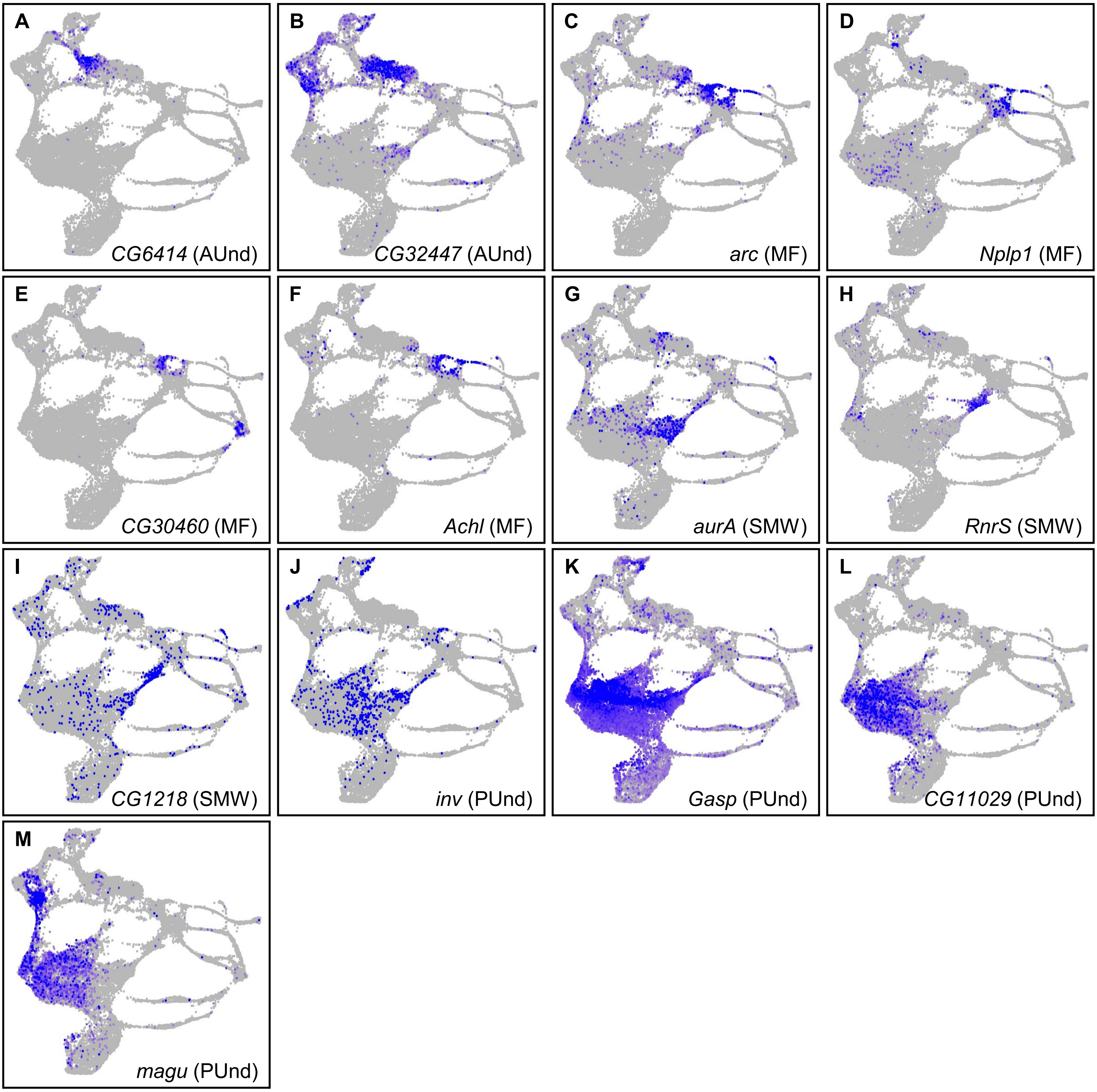

A
*CG6414* (AUnd)
B
*CG32447* (AUnd)
C
*arc* (MF)
D
*Nplp1* (MF)
E
*CG30460* (MF)
F
*Achl* (MF)
G
*aurA* (SMW)
H
*RnrS* (SMW)
I
*CG1218* (SMW)
J
*inv* (PUnd)
K
*Gasp* (PUnd)
L
*CG11029* (PUnd)
M
*magu* (PUnd)

**Supplementary Fig. 4: Novel markers of undifferentiated cell clusters.** *CG6414* (**A**), and *CG32447* (**B**) are novel AUnd markers. *arc* (**C**), *Neuropeptide-like precursor 1* (*Nplp1*, **D**), *CG30460* (**E**), and *Achilles* (*Achl*, **F**) are primarily expressed in the MF. *aurora A* (*aurA*, **G**), *Ribonucleoside diphosphate reductase small subunit* (*RnrS*, **H**), and *CG1218* (**I**) are three novel SMW markers. **K-N.** Novel PUnd markers: *invected* (*inv*, **J**), *Gasp* (**K**), *CG11029* (**L**), and *magu* (**M**).

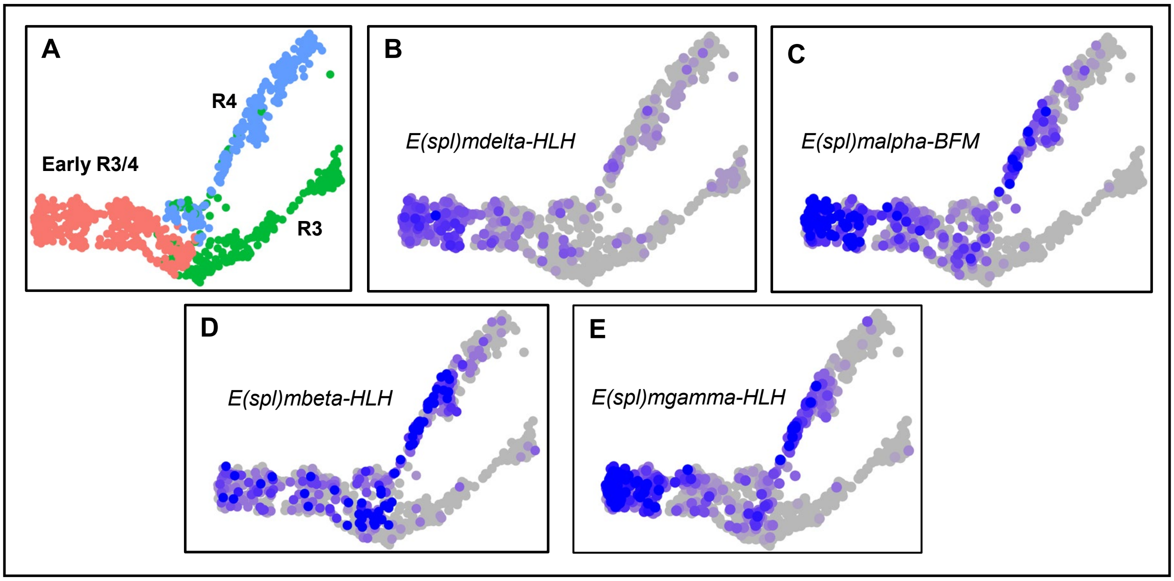
A
R4
Early R3/4
R3
B
E(spl)mdelta-HLH
C
E(spl)malpha-BFM
D
E(spl)mbeta-HLH
E
E(spl)mgamma-HLH

**Supplementary Fig. 5: The R4 cell cluster shows higher levels of *Enhancer of split* gene expression. A.** UMAP plot showing the split in the R3/4 stream. **B-E.** FeaturePlots showing the expression of several *E(spl)* genes: **B.** *E(spl)mdelta-HLH*, **C.** *E(spl)malpha-BFM*, **D.** *E(spl)mbeta-HLH*, and **E.** *E(spl)mgammaa-HLH*. *E(spl)* genes show higher levels of expression in R4 compared to R3.

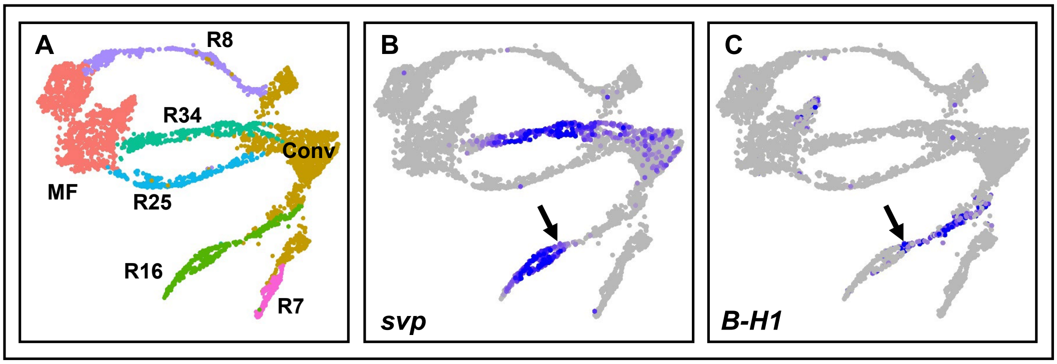
A
R8
R34
Conv
MF
R25
R16
R7
B
svp
C
B-H1

**Supplementary Fig. 6: Photoreceptor streams comprise cells in a temporal developmental series.**

**A.** UMAP plot showing the MF and PRs only. PR subtype clusters appear as streams emerging from the MF (R8, R2/5, and R3/4) or undifferentiated cells (R1/6 and R7). **B.** A FeaturePlot showing the expression of *svp* thoughout the R3/4 stream but only in less mature R1/6 cells. **C.** *B-H1* FeaturePlot showing expression in mature but not developmentally younger R1/6 cells. *B-H1* mRNA is observed only after *svp* expression ceases along the stream.

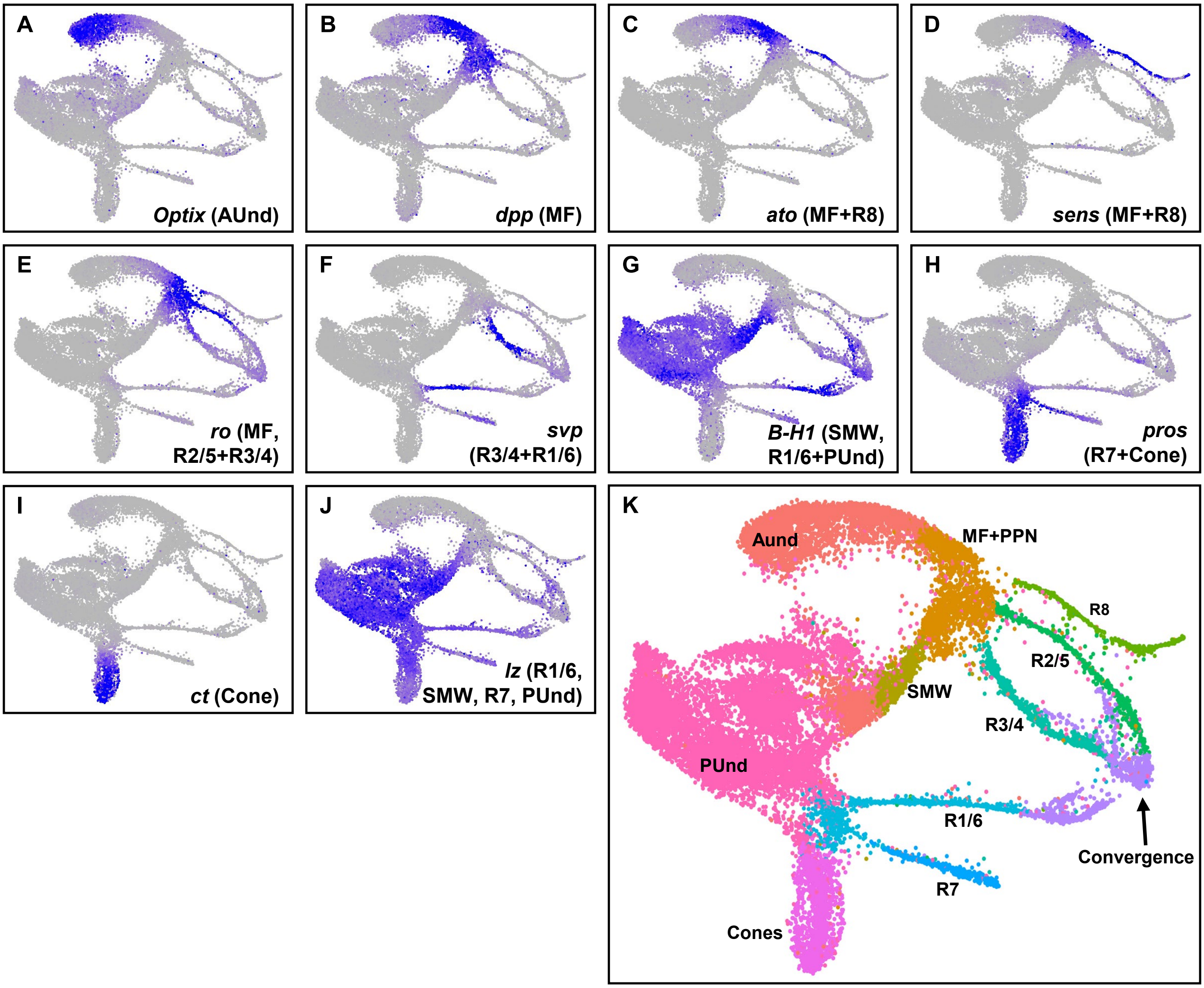
A
Optix (AUnd)
B
dpp (MF)
C
ato (MF+R8)
D
sens (MF+R8)
E
ro (MF, R2/5+R3/4)
F
svp (R3/4+R1/6)
G
B-H1 (SMW, R1/6+PUnd)
H
pros (R7+Cone)
I
ct (Cone)
J
lz (R1/6, SMW, R7, PUnd)
K
Aund
MF+PPN
R8
R2/5
SMW
R3/4
PUnd
R1/6
Convergence
R7
Cones

**Supplementary Fig. 7: Annotation of snATAC-seq cell clusters by imputation of RNA from scRNA-seq data. A-J.** FeaturePlots showing the distribution of imputed RNA on snATAC-seq clusters. **A.** FeaturePlot showing *Optix* mRNA in the AUnd cluster. **B.** *decapentaplegic* (*dpp*) mRNA is predominantly detected in the MF. **C.** *atonal* (*ato*) mRNA is in the MF and R8 stream. **D.** *senseless* (*sens*) mRNA is specific to the MF and R8. **E.** *rough* (*ro*) mRNA is confined to the MF, R2/5 and R3/4 cell clusters. **F.** *seven up* (*svp*) mRNA is detected primarily in R3/4 and R1/6. **G.** *B-H1* mRNA is observed in R1/6 and PUnd. **H.** *prospero* (*pros*) is specifically present in the R7 and cone cell clusters. **I.** *cut* (*ct*) is completely confined to cone cells. **J.** FeaturePlot of *lozenge* (*lz*) showing expression primarily in R1/6, R7 and PUnd. **K.** The UMAP plot for late larval snATAC-seq data with cluster identities is shown for reference.

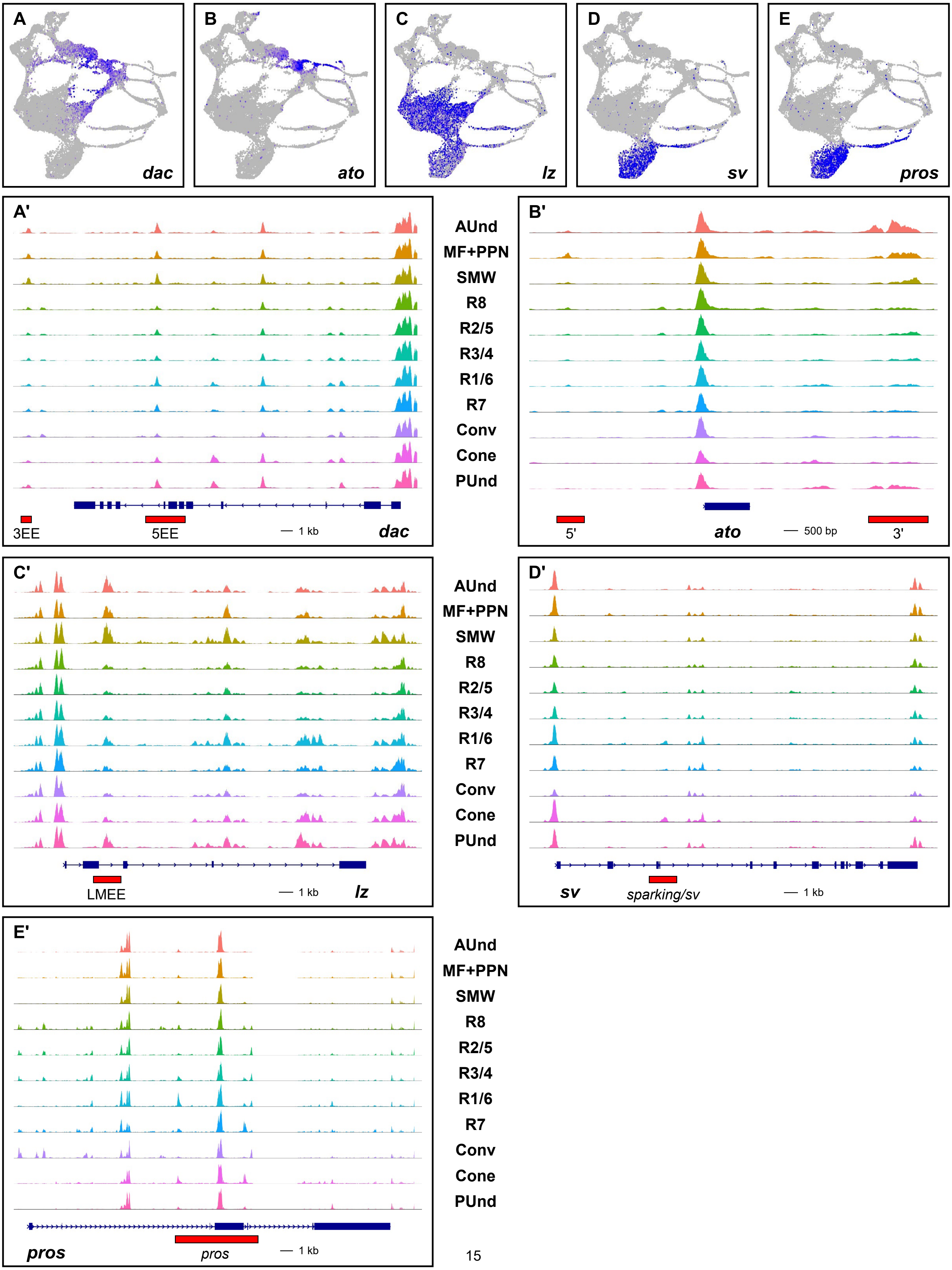

A
dac
B
ato
C
lz
D
sv
E
pros
A'
AUnd
MF+PPN
SMW
R8
R2/5
R3/4
R1/6
R7
Conv
Cone
PUnd
3EE
5EE
1 kb
dac
B'
5'
ato
500 bp
3'
C'
LMEE
1 kb
lz
D'
sv
sparking/sv
1 kb
E'
pros
pros
1 kb

**Supplementary Fig. 8: Validation of snATAC-seq clusters using known larval eye enhancers. A-E.** FeaturePlots showing the expression of the known marker genes *dachshund* (*dac*, **A**), *ato* (**B**), *lz* (**C**), *shaven* (*sv*, **D**), and *pros* (**E**). **A'-E'** show snATAC-seq CoveragePlots of known enhancer regions corresponding to the genes shown in **A-E.** The red horizontal box in each CoveragePlot denotes the relative size and position of known enhancer sequences that are sufficient to drive reporter gene expression in the eye. **A'**. *dac* CoveragePlot showing the 3EE and 5EE enhancers. The peak corresponding to 3EE is specifically accessible in the AUnd, MF+PPN, and SMW cell clusters. **B'.** The 5' and 3' enhancers of *ato* span regions of accessible chromatin. **C'.** CoveragePlot of *lz* showing a cone-specific peak (albeit somewhat weak) in the same position as the LMEE enhancer. **D'**. CoveragePlot of *sv* showing a cone-specific peak. The *sparkling/sv* enhancer maps to this peak region. **E'**. *pros* CoveragePlot showing an R7-enriched peak that is within the *pros* enhancer.

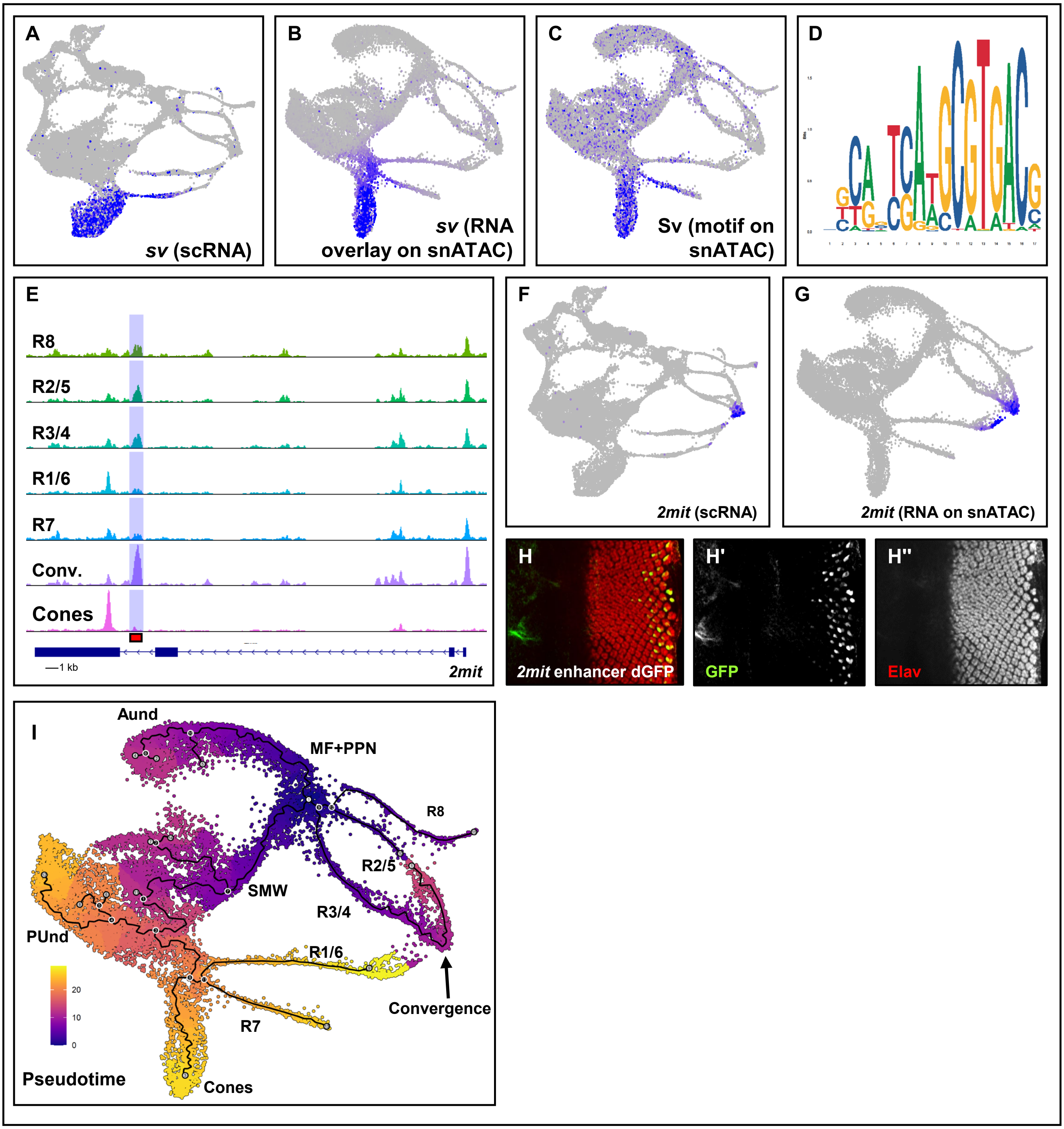
A
sv (scRNA)
B
sv (RNA overlay on snATAC)
C
Sv (motif on snATAC)
D
E
R8
R2/5
R3/4
R1/6
R7
Conv.
Cones
1 kb
2mit
F
2mit (scRNA)
G
2mit (RNA on snATAC)
H
2mit enhancer dGFP
H'
GFP
H''
Elav
I
Aund
MF+PPN
R8
R2/5
SMW
R3/4
PUnd
R1/6
R7
Convergence
Cones
20
10
0
Pseudotime

**Supplementary Fig. 9: Validation of snATAC-seq clusters using motif analyses and identification of novel enhancers. A.** scRNA-seq FeaturePlot of *sv* showing expression in R7 and cone cell clusters. **B.** FeaturePlot with the distribution of imputed *sv* RNA on snATAC-seq clusters showing RNA in R7 and cones. **C.** FeaturePlot showing the distribution of the Sv motif on the snATAC-seq UMAP plot. The Sv motif is enriched in R7 and cone cells. **D.** A Sv motif logo showing the Sv binding site. **E.** CoveragePlot of *2mit* showing a Convergence-enriched peak highlighted in blue. The red bar indicates the DNA sequence used to make a reporter transgene. **F.** FeaturePlot of *2mit* showing expression mRNA expression in the Convergence cell cluster. **G.** *2mit* RNA overlaid on the snATAC-seq UMAP plot showing Convergence-specific distribution of mRNA. **H-H''.** Transgenic larval eye disc carrying a transgene with *2mit* peak-DNA driving destabilized GFP. The eye disc is costained with GFP (green) and Elav (red). GFP is detected in the most posterior ommatidial columns (**H** and **H'**). **I.** snATAC-seq UMAP plot showing pseudotime trajectories inferred by Monocle 3. R1/6, R7 and cones show late pseudotime. The early pseudotime is indicated by purple, whereas yellow color represents a late pseudotime.

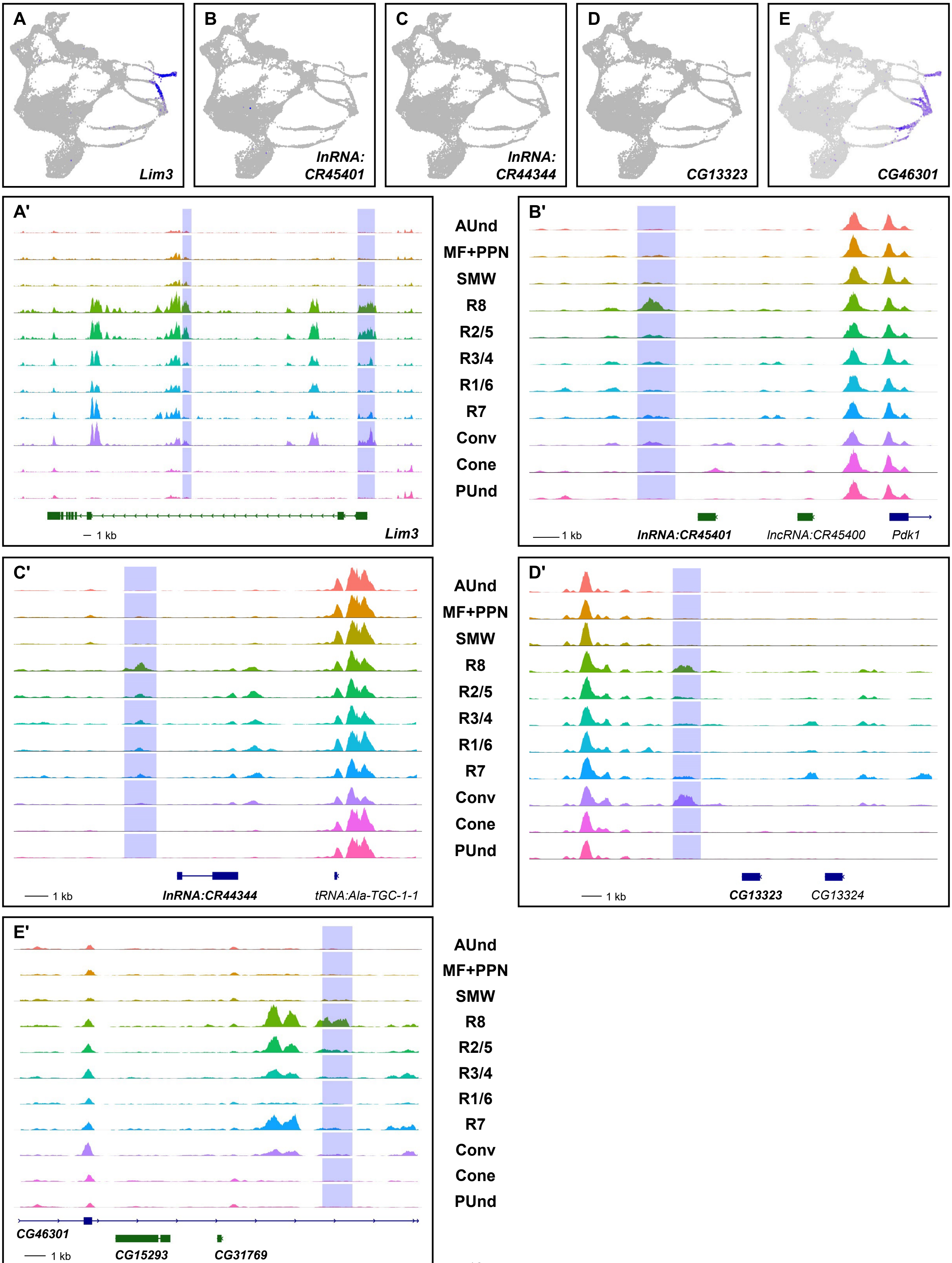
A
Lim3
B
lnRNA: CR45401
C
lnRNA: CR44344
D
CG13323
E
CG46301
A'
AUnd
MF+PPN
SMW
R8
R2/5
R3/4
R1/6
R7
Conv
Cone
PUnd
1 kb
Lim3
B'
1 kb
lnRNA:CR45401
lncRNA:CR45400
Pdk1
C'
1 kb
lnRNA:CR44344
tRNA:Ala-TGC-1-1
D'
1 kb
CG13323
CG13324
E'
CG46301
1 kb
CG15293
CG31769

**Supplementary Fig. 10: Differentially accessible R8 cluster-specific snATAC-seq peaks.**

**A-E.** FeaturePlots showing the expression of *Lim3* (**A**), *lnRNA:CR45401* (**B**), *lnRNA:CR44344* (**C**), *CG13323* (**D**), and *CG31769* (**E**). *Lim3* is expressed in R8 and R2/5. No mRNA is detected on the FeaturePlots of the other genes. **A'-E'.** CoveragePlots corresponding to the genes shown in **A-E.** All CoveragePlots show peaks that are primarily R8-specific.

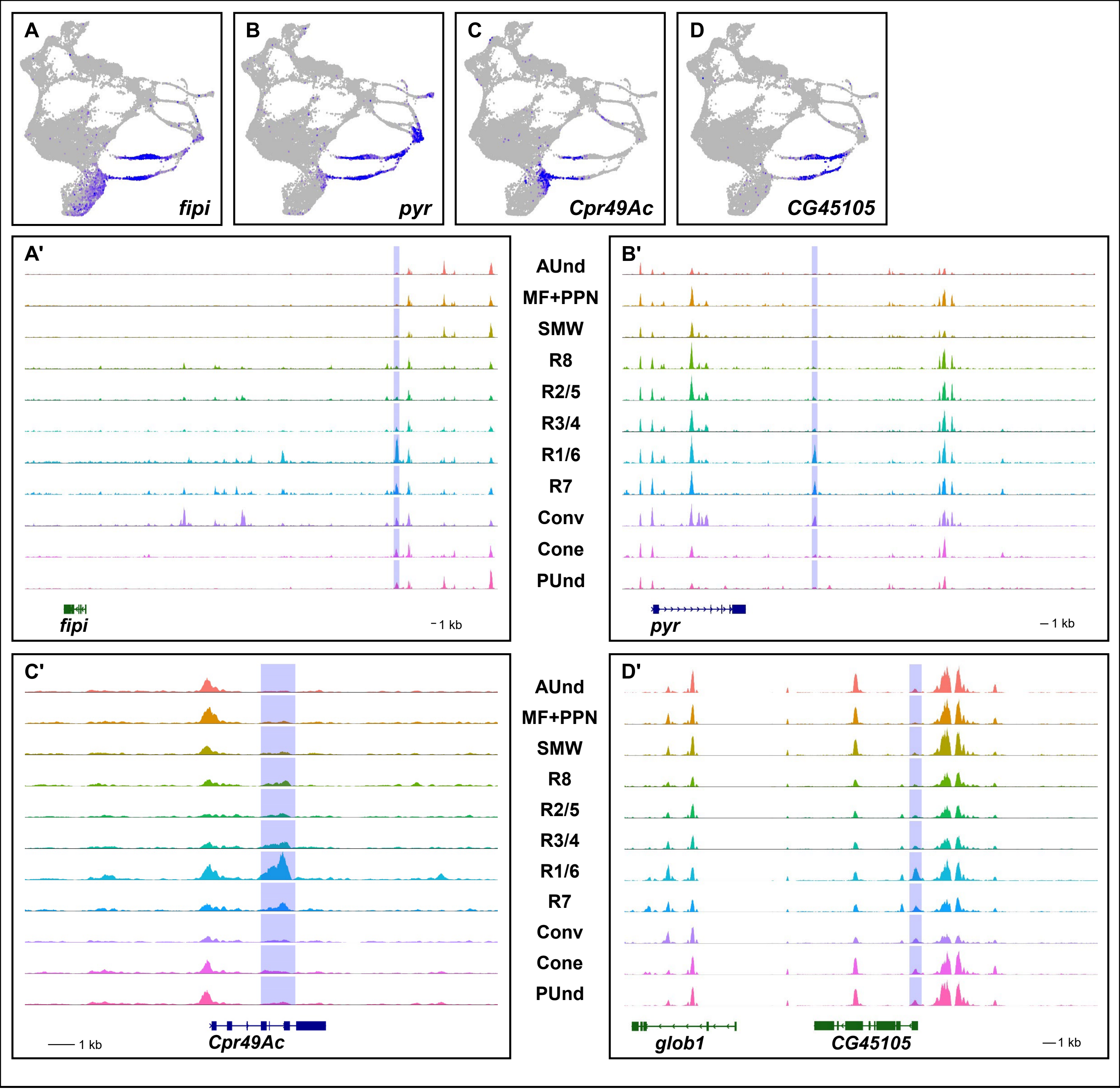
A
fipi
B
pyr
C
Cpr49Ac
D
CG45105
A'
B'
AUnd
MF+PPN
SMW
R8
R2/5
R3/4
R1/6
R7
Conv
Cone
PUnd
fipi
1 kb
pyr
1 kb
C'
D'
Cpr49Ac
1 kb
glob1
CG45105
1 kb

**Supplementary Fig. 11: Differentially accessible R1/6 and R7 cluster-specific peaks. A-D.** FeaturePlots showing the expression of *fipi* (**A**), *pyr* (**B**), *Cpr49Ac* (**C**), and *CG45105* (**D**). *fipi*, *pyr*, *Cpr49Ac*, and *CG45105* are expressed in both R1/6 and R7. *fipi* and *Cpr49Ac* are also expressed in the cone cell cluster while *pyr* is also preesent in the Convergence cluster. **A'-D'.** CoveragePlots corresponding to the genes shown in **A-D**. All CoveragePlots show peaks largely specific to the R1/6 and R7 cell clusters.

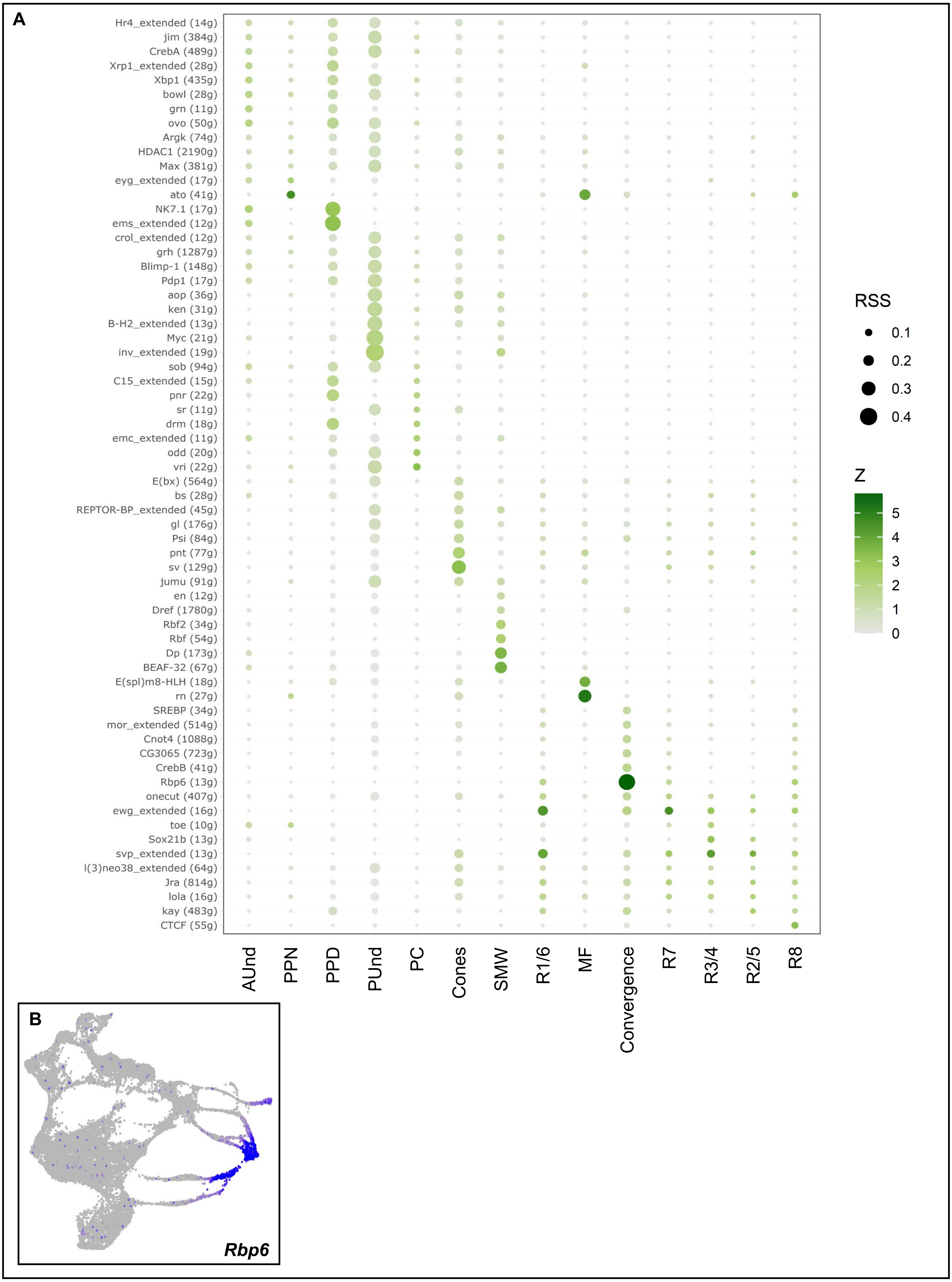
A
Hr4_extended (14g)
jim (384g)
CrebA (489g)
Xrp1_extended (28g)
Xbp1 (435g)
bowl (28g)
grn (11g)
ovo (50g)
Argk (74g)
HDAC1 (2190g)
Max (381g)
eyg_extended (17g)
ato (41g)
NK7.1 (17g)
ems_extended (12g)
crol_extended (12g)
grh (1287g)
Blimp-1 (148g)
Pdp1 (17g)
aop (36g)
ken (31g)
B-H2_extended (13g)
Myc (21g)
inv_extended (19g)
sob (94g)
C15_extended (15g)
pnr (22g)
sr (11g)
drm (18g)
emc_extended (11g)
odd (20g)
vri (22g)
E(bx) (564g)
bs (28g)
REPTOR-BP_extended (45g)
gl (176g)
Psi (84g)
pnt (77g)
sv (129g)
jumu (91g)
en (12g)
Dref (1780g)
Rbf2 (34g)
Rbf (54g)
Dp (173g)
BEAF-32 (67g)
E(spl)m8-HLH (18g)
rn (27g)
SREBP (34g)
mor_extended (514g)
Cnot4 (1088g)
CG3065 (723g)
CrebB (41g)
Rbp6 (13g)
onecut (407g)
ewg_extended (16g)
toe (10g)
Sox21b (13g)
svp_extended (13g)
l(3)neo38_extended (64g)
Jra (814g)
lola (16g)
kay (483g)
CTCF (55g)
AUnd
PPN
PPD
PUnd
PC
Cones
SMW
R1/6
MF
Convergence
R7
R3/4
R2/5
R8
RSS
0.1
0.2
0.3
0.4
Z
5
4
3
2
1
0
B
Rbp6

**Supplementary Fig. 12: Top regulons of each scRNA-seq cell cluster identified using SCENIC. A.** RSS plot showing the top regulons in each cell cluster. The color intensity of each dot is proportional to the expression level of each regulon. The size of the dot is proportional to the regulon specificity score. **B.** FeaturePlot showing the expression of *Rbp6* in the Convergence cluster. *Rbp6* is highly expressed and specific to the Convergence cell cluster and late R1/6 cells.

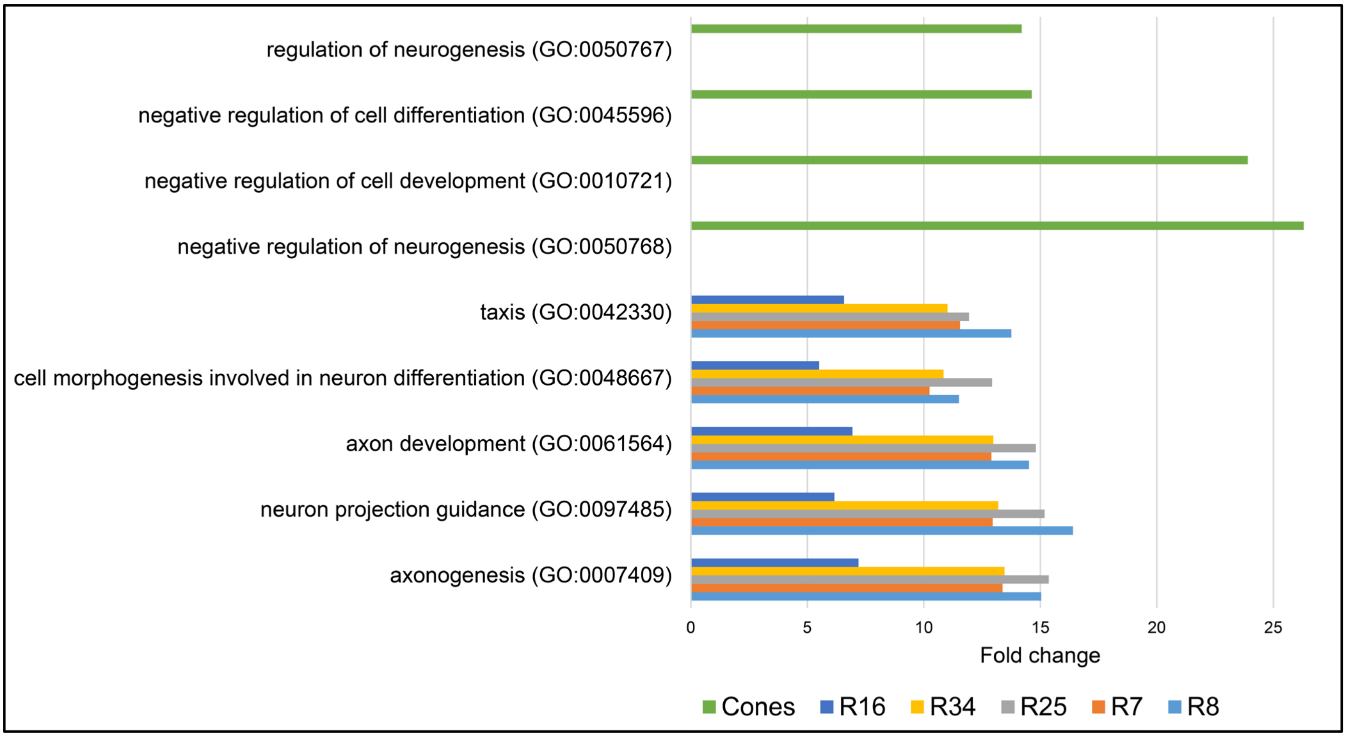
regulation of neurogenesis (GO:0050767)
negative regulation of cell differentiation (GO:0045596)
negative regulation of cell development (GO:0010721)
negative regulation of neurogenesis (GO:0050768)
taxis (GO:0042330)
cell morphogenesis involved in neuron differentiation (GO:0048667)
axon development (GO:0061564)
neuron projection guidance (GO:0097485)
axonogenesis (GO:0007409)
0
5
10
15
20
25
Fold change
Cones
R16
R34
R25
R7
R8

**Supplementary Fig. 13: Top enriched Gene Ontology terms for PR and cone cell marker peaks.** Bar graphs showing the fold enrichment of the top enriched Gene Ontology (GO) terms for genes that are near differentially accessible PR peaks. GO terms related to axon development are enriched in PR clusters, while cone cells show enrichment for negative regulation of neurogenesis.
